# Altitude-dependent agro-ecologies impact the microbiome diversity of scavenging indigenous chicken in Ethiopia

**DOI:** 10.1101/2023.06.12.544316

**Authors:** Laura Glendinning, Xinzheng Jia, Adebabay Kebede, Samuel O. Oyola, Jong-Eun Park, Woncheoul Park, Abdulwahab Assiri, Jacob Bak Holm, Karsten Kristiansen, Jianlin Han, Olivier Hanotte

## Abstract

Scavenging indigenous village chickens play a vital role in sub-Saharan Africa, sustaining the livelihood of millions of farmers. These chickens are exposed to vastly different environments and feeds compared to commercial chickens. In this study, we analysed the caecal microbiota of 243 Ethiopian village chickens living in different altitude-dependent agro-ecologies. Differences in bacterial diversity were significantly correlated with differences in specific climate factors, topsoil characteristics, and supplemental diets provided by farmers. Microbiota clustered into 3 enterotypes, with one particularly enriched at high altitudes. We assembled 9,977 taxonomically and functionally diverse metagenome-assembled genomes, the vast majority of which were not found in a dataset of previously published chicken microbes, or in the Genome Taxonomy Database. The wide functional and taxonomic diversity of these microbes highlights their importance in the local adaptation of indigenous poultry, and the significant impacts of environmental factors on the microbiota argues for further discoveries in other agro-ecologies.

## Introduction

Scavenging or semi-scavenging indigenous village chickens play a key role in sub-Saharan African and Asian countries, sustaining the livelihood of millions of farmers. They predominantly comprise indigenous chicken genotypes well adapted to the local environment^1^. As these birds are exposed to high predation and diseases challenges^2^, survival traits rather than production traits are favoured by natural selection, and to some extent, human selection. These chickens also contribute to the spread of zoonotic diseases such as campylobacteriosis and salmonellosis, which have a high disease burden in African populations^3, 4^. By understanding the microbiota of these chickens, we may be able to suggest ways of improving their nutrition and disease resistance, and henceforth productivity.

Over 95% of poultry products sold in Ethiopia originate from indigenous village chickens^5^. Ethiopia is an ecologically diverse country with distinct environmental zones, ranging from the hot and arid climate of the lowlands to the cold and humid climate of the highlands. These climatic zones, and a diverse geographical topography, form naturally varied environmental conditions for smallholder crop-livestock farming systems. Geographical location^6^, temperature^7^, and altitude^8^ have previously been demonstrated to impact the composition of the chicken microbiota using 16S rRNA gene amplicon sequencing.

The gut microbiota plays important roles in chicken health and productivity, contributing to nutrition, immune development, and pathogen resistance^9^. The highest concentration of microbes in chicken can be found in the caeca. Here the microbial communities play a vital nutritional role, fermenting fibre into short-chain fatty acids (SCFAs) that can be used as an energy source by the bird, as well as contributing to colonisation resistance, and nitrogen recycling^10, 11^.

Through the use of culturing techniques^12, 13^, metabarcoding^14, 15^ and metagenomics^13, 16, 17^, there have been many recent advances in our understanding of the chicken caecal microbial ecosystems. However, most of these studies have examined grain-fed, commercial chicken breeds that were hatched and housed in biosecure facilities without maternal contact. The target of these commercial-like conditions is to enhance productivity by standardising/controlling host-environment interactions. Commercial chicken breeds, and by extension their microbiota, have also been shaped by human selection for high productivity traits. On the contrary, scavenging or semi-scavenging chicken population are exposed to far greater predation and diseases challenges^2^, and therefore survival traits have been selected for rather than production. Not only has this affected the chicken genome^1^, but would also reasonably be expected to have impacted the gut microbiota.

To characterise the microbiota of indigenous chickens in Ethiopia, we collected 243 chicken caecal content samples from 26 villages in 15 districts. Shotgun metagenomics was used to characterise the caecal microbiota. In order to profile all taxa, including low-abundance taxa, we constructed a catalogue of non-redundant genes. We identified three distinct enterotypes, one of which was particularly elevated in the highest altitude samples. We also constructed metagenome-assembled genomes (MAGs). Previously MAGs have been constructed from chicken breeds including Ross 308s, Lohman Browns and Silkies^13, 16, 17^. We constructed 9,977 high-quality, strain level MAGs and 1,790 species-level MAGs, representing diverse taxonomies. We found that the vast majority of the MAGs were not present in a dataset of microbial genomes from previous chicken microbiota studies. The MAGs generated in this study contained genes encoding a large diversity of carbohydrate active enzymes (CAZymes) and metabolic functions.

## Results

After the removal of three samples during quality control, we characterised the microbiota of the caecal contents of 240 indigenous Ethiopian scavenging chickens from 26 sampling sites, aiming to further understand the impact of agro-ecology on the gut microbiota composition. The sampling sites were highly diverse, representing different latitudes, longitudes and altitudes (**Figure 1B**). Five distinct climate zones were defined based on climate variation analysis, with altitude and annual mean temperature as major predictors (**Figure 1C**).

**Figure 1:**
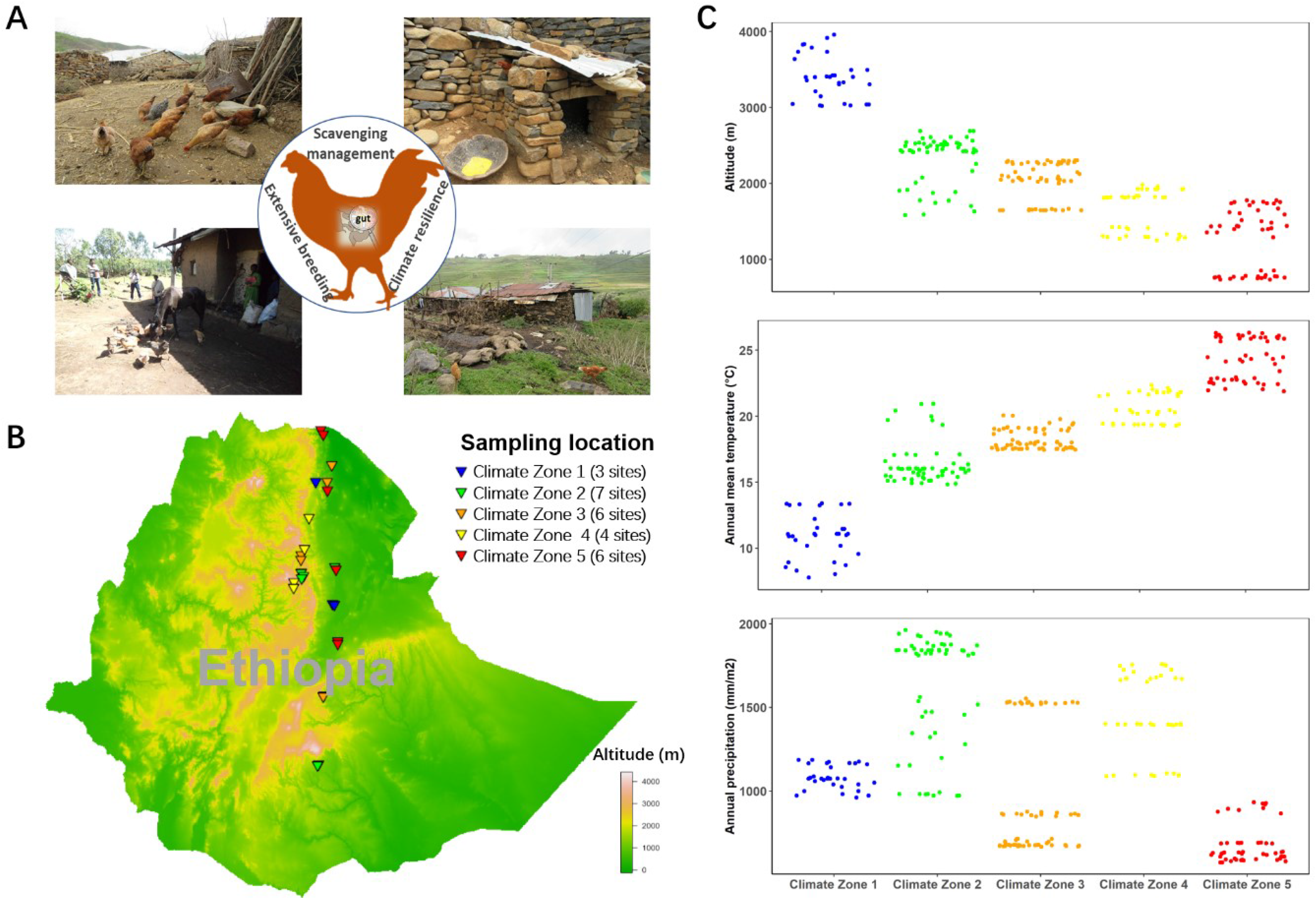
Sample site information for Ethiopian indigenous chickens under scavenging production systems. A) Examples of scavenging production systems in Ethiopia. B) Geographic distribution of sampling sites within Ethiopia. C) Five climate zones were defined based on climate factors. These included altitude, precipitation, and temperature, as shown.

### Construction of an Ethiopian indigenous chicken microbiota gene catalogue

By constructing a gene catalogue we can detect more rare taxa than by using MAGs alone, and also identify taxa from which it is difficult to construct high-quality MAGs due to their large genome sizes or complex genome structures. We constructed a reference gene catalogue of 33,629,587 non-redundant genes (**Figure S1**). Rarefaction analysis of sampling size revealed that 90% of genes were captured with a sampling size of 60 individuals (**Figure S2**). Genes that are identified across many individuals within a population can be defined as “core genes”. Only 420,891 (1.2%) genes were shared amongst 80% of samples. This may be due to the high inter-individual diversity of microbiota-derived genes within the caecal samples, or due to insufficient sequencing depth to detect total gene numbers.

We next characterised the taxonomic origin of the genes in the gene catalogue. Fifty-five percent (55%) of the genes were assigned a taxonomic label. Of these genes, 87.9%, 56.5% and 49.2% were assigned to a phylum, genus, and species, respectively (**Figure 2A**). After removing DNA that likely originated from the host diet (plants and insects), a total of 33,435,297 genes remained that were used for microbiota analysis.

**Figure 2:**
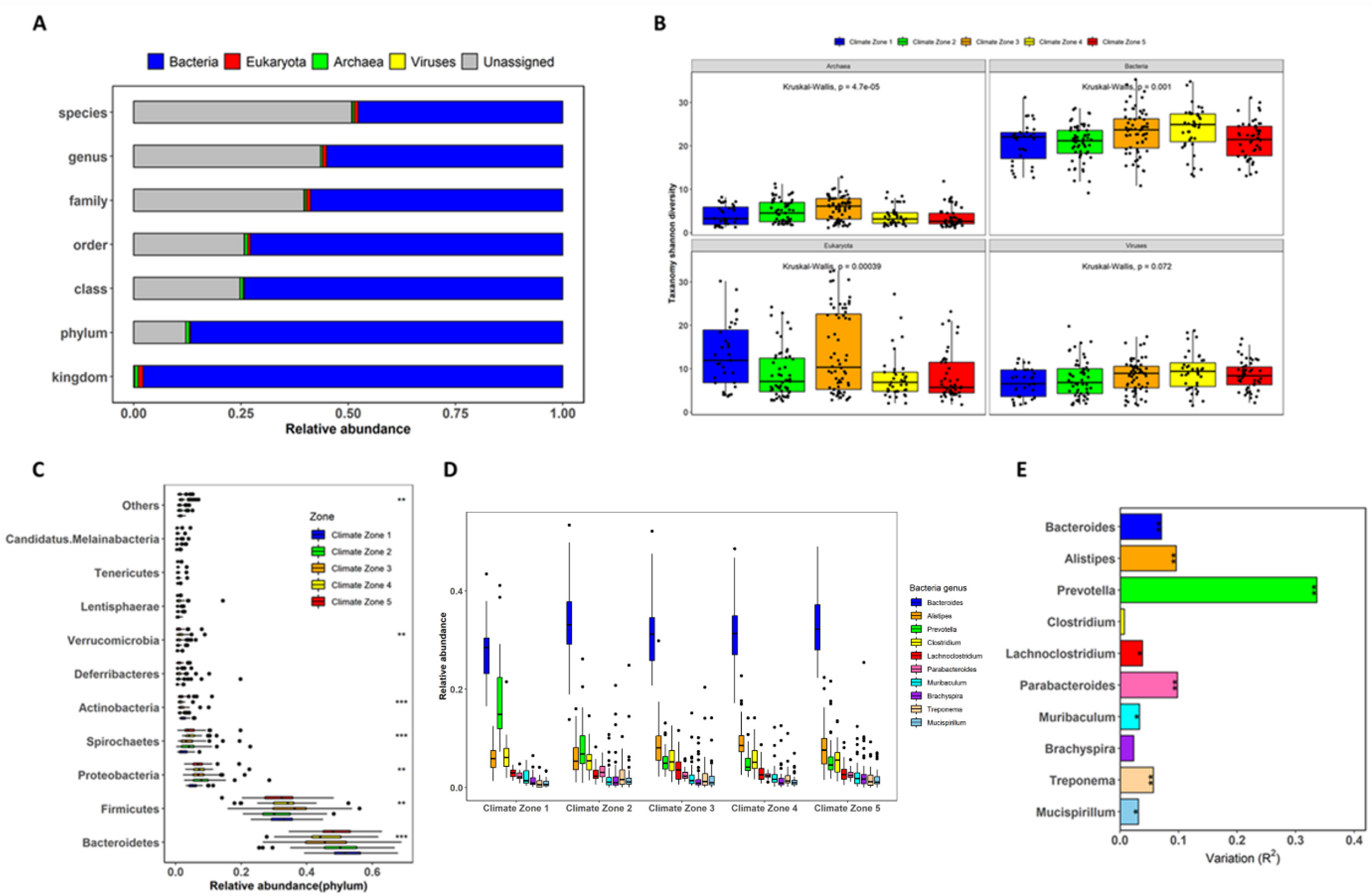
Structure and phylogenetic diversity of the caecal microbiota from indigenous Ethiopian chickens. A) Abundance distribution of taxa at each phylogenetic level B) Alpha-diversity (Shannon index) of the archaea, bacteria, viruses and Eukaryota in caecal samples. C) Relative abundance of microbiota at phylum level among climate zones (** p < 0.01; *** p < 0.001). D) Genus relative abundance among climate zones. E) Comparison of genera abundance among populations from distinct climate zones.

We focused our analysis on taxa averaging at least one count per million sequences. Each sample had an average relative abundance of 98% bacteria (± 1.22%), 0.86% archaea (± 0.54%), 0.98% Eukaryota (± 1.08%) and 0.16% viruses (± 0.13%). Bacteroidota represented the most abundant phylum (48.0%), followed by Firmicutes (32.9%), Proteobacteria (7.3%), Spirochaetota (4.2%), Actinobacteriota (1.9%) and Deferribacterota (1.1%). Six of the 10 most abundant phyla showed significant differences in abundance between climate zones (**Figure 2C**). The most abundant genera were *Bacteroides* (31.9%), *Alistipes* (7.9%) and *Prevotella* (7.6%), all members of the Bacteroidota phylum. *Prevotella,* in particular, showed a high level of variation between climate zones (**Figure 2D-E**). Notably, chickens from climate zone 1 (high altitude) had more than 2.5-fold higher abundance of *Prevotella* (17.6%) than chickens from other climate zones. Archaea, bacteria, and Eukaryota showed significant differences in alpha-diversity between climate zones (Kruskal-Wallis p value < 0.01), while viruses showed no significant differences (**Figure 2B**). For bacteria, diversity gradually increased from climate zone 1 (> 3000 m – high altitude) to climate zone 4 (around 1300 m – medium altitude), and was slightly decreased in climate zone 5 (around 1000 m).

### Indigenous Ethiopian chickens contain three enterotypes related to environmental distribution

Using genera abundances as estimated from the gene catalogue, the Ethiopian chicken caecal microbiota clustered into three enterotypes (**Figure 3A**). Samples belonging to enterotype 3 were particularly distinct from enterotypes 1 and 2. Climate zone 1 is clearly dominated by microbiota of enterotype 3, which accounts for 62% of samples from this zone (**Figure 3B**). Linear discriminant analysis Effect Size (LEfSe) analysis was carried out to identify differential enrichment of genera between enterotypes. For enterotype 1, the top ten most discriminating genera were *Alistipes*, *Treponema*, *Brachyspira*, *Mucispirillum*, *Muribaculum*, *Parabacteroides*, *Sutterella*, *Acidaminococcus*, *Sphaerochaeta* and *Akkermansia*; for enterotype 2 the top genera were *Bacteroides*, *Lachnoclostridium*, *Clostridium*, *Blautia*, *Pseudoflavonifractor*, *Flavonifractor*, *Erysipelatoclostridium*, *Azospirillum*, *Merdimonas* and *Gemmiger*; and for enterotype 3 *Prevotella*, *Megamonas*, *Faecalibacterium*, *Olsenella*, *Lactobacillus*, *Bifidobacterium*, *Mediterranea*, *Collinsella*, *Megasphaera* and *Dialister* (**Figure 3C and Figure 3D**). Co-occurrence networks based on correlation analysis revealed that the abundance of major discriminating genera had strong positive correlations with the abundance of other genera (**Figure 3E**).

**Figure 3:**
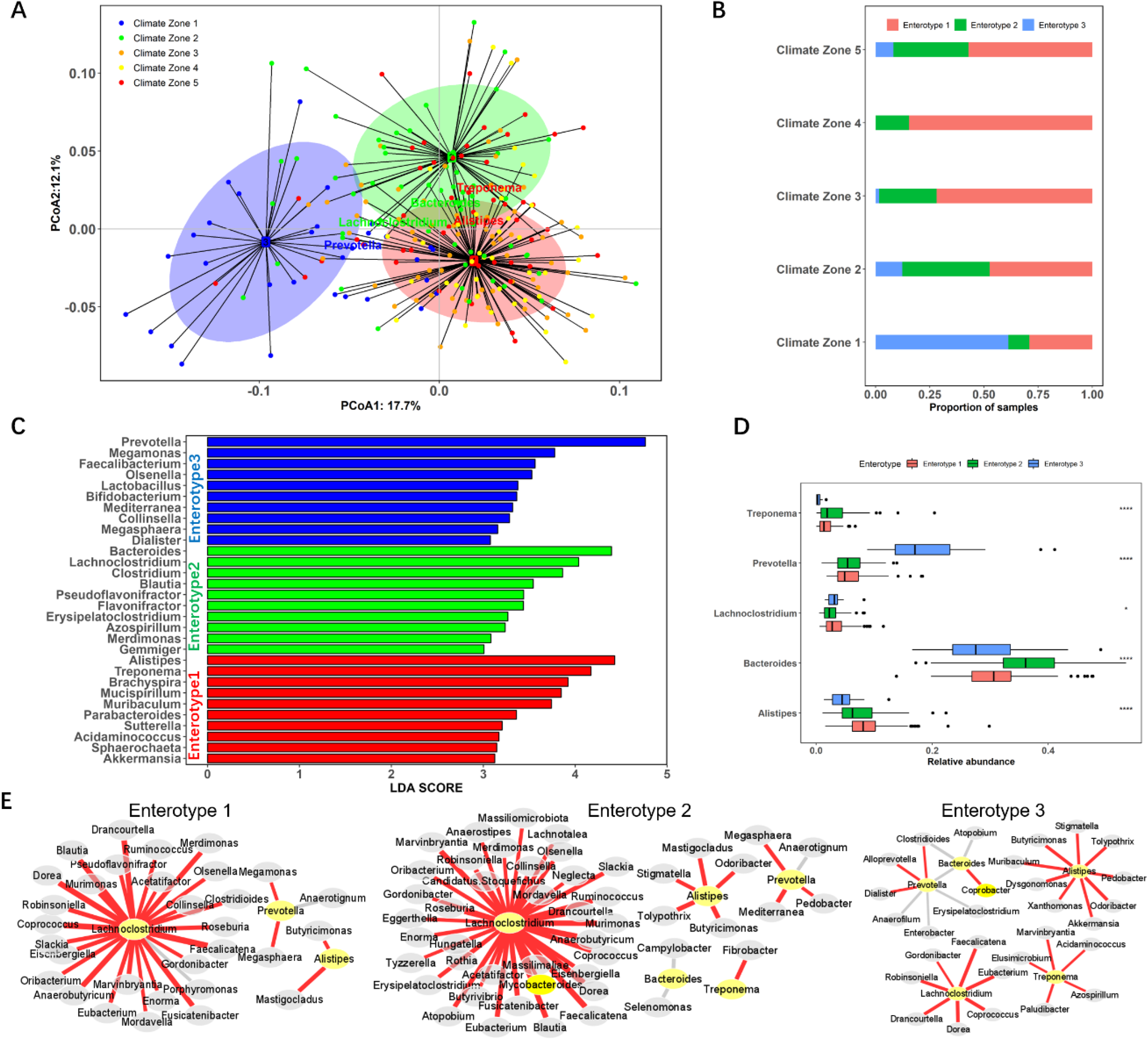
Three enterotypes observed in the caecal microbiota of Ethiopian indigenous chickens. A) Caecal microbiota enterotypes clustered by PCA, with discriminating genera highlighted. B) Distribution of enterotypes as proportions of samples from different climate zones. C) Top ten highest linear discriminant analysis (LDA) scores for genera contributing to the discrimination of each enterotype. D) Abundance profiles of the main genera contributing to each enterotype, as defined by LEfSe (* p < 0.05; *** p < 0.001). E) Co-occurrence networks based on correlations between genera abundance. Red and grey lines indicated positive and negative correlations (p < 0.01, rho > 0.5 relative abundance > 0.01%). Yellow nodes represented the main genus contributors (LDA score > 4).

### Agro-ecological factors contributing to microbiota composition

We next wanted to identify which agro-ecological factors were correlated with differences in the beta-diversity of the caecal microbiota, using the gene catalogue data. Beta-diversity was significantly associated with supplementary diets provided by farmers and the location’s topsoil characteristics. Significant differences in beta-diversity (Bray-Curtis dissimilarity: genera, species and strains) were associated with temperature, altitude, precipitation and seasonal cycles, followed by supplementary diets, co-raising styles and local topsoil contents (adj-p < 0.01). Redundancy analysis was used to capture the driving factors contributing to microbiota diversity. For climate factors: altitude and seasonal precipitation (bio 15) were the major factors, explaining 10% of the total variation (**Figure 4A**). Cation exchange capacity of topsoil (CECSOL) and silt percentage of topsoil (SLTPPT) were identified as major contributors to diversity, explaining 11% of variation (**Figure 4B**). The provision of supplementary diets to chickens was also considered. We found that three common grains (maize, wheat and barley) significantly impacted the microbiota diversity, explaining 8% of total variation. Microbiota diversity significantly increased as altitude increased (P < 0.01, **Figure 4C**). Amongst the main taxonomic contributors to enterotypes, *Prevotella* and *Megamonas* were significantly positively correlated to altitude (**Figure 4D and 4E**), while *Corallococcus* was negatively correlated (**Figure 4F**).

**Figure 4:**
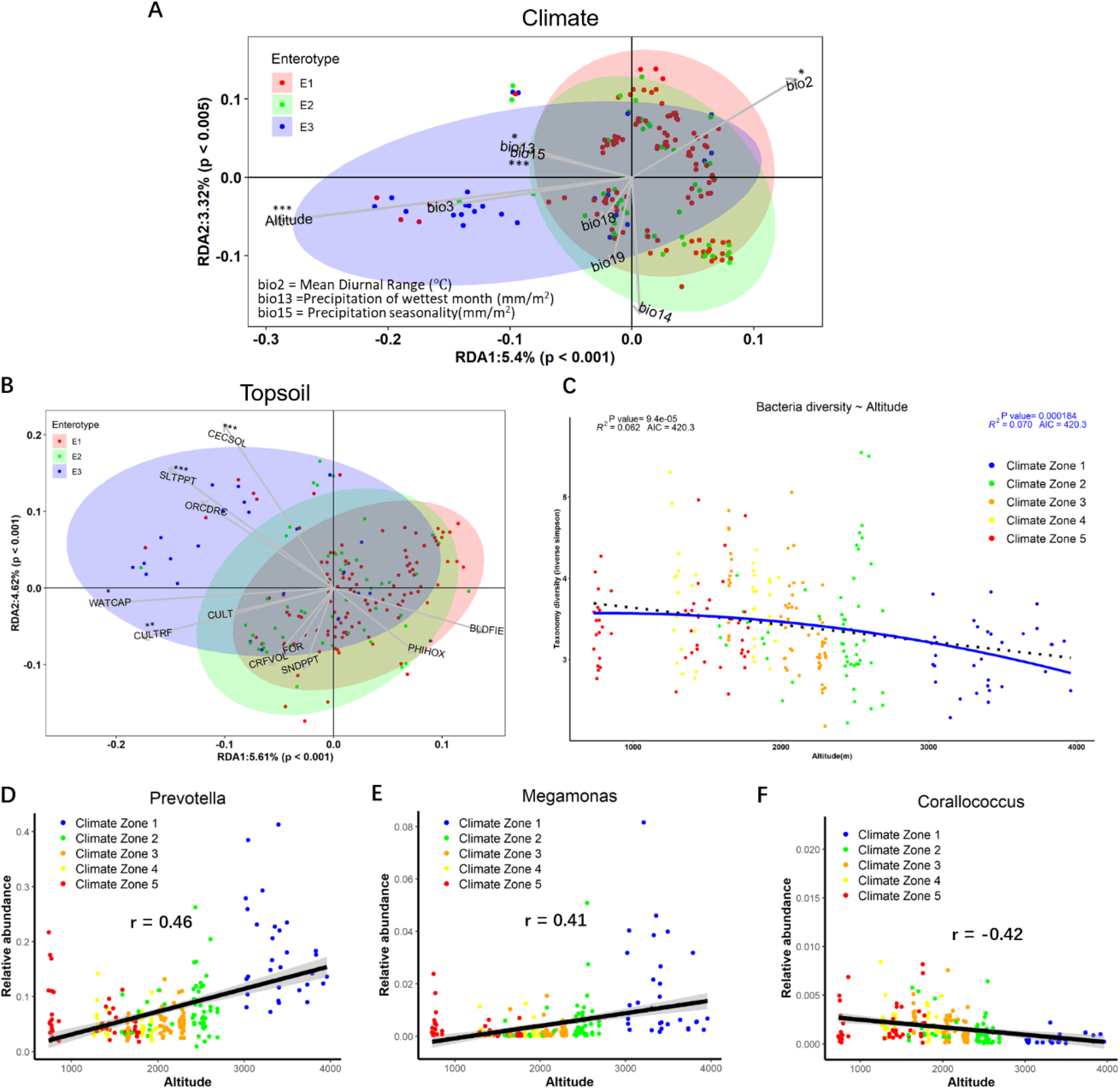
Microbiota composition exhibited significant geographic and ecological diversity. A) RDA analysis on the microbiota composition of samples (gene catalogue) and climate factors including altitude, temperature and precipitation. B) RDA analysis on the bacteria composition and ecological factors including common topsoil characteristics. C) Bacterial diversity (Inverse Simpsons index) decreases with altitude gradient. The best polynomial fit in blue was determined on the basis of the corrected Akaike Information Criterion (AIC) of the order polynomial models. The ANOVA test was used to test for significance. D-F) Spearman correlation was performed to test the relationship between the abundance of genera and altitude.

### Assembly of 9,977 microbial genomes from diverse taxonomies

Whilst gene catalogues can provide information about the taxonomies present in a dataset, including those that are present only in low abundances, it is also possible to generate genomic level information about components of the microbiota by constructing metagenome assembled genomes (MAGs) from the more abundant taxa in the samples.

We constructed 9,977 high-quality, non-redundant MAGs at strain level, and 1,790 at species level (**Figure S3**, **Supplementary table 1**). For strain-level MAGs, 9815 were identified as bacteria and 162 were identified as Archaea (**Supplementary table 2**). Archaea comprised 3 phyla, Halobacteriota (n = 39), Methanobacteriota (n = 28) and Thermoplasmatota (n = 95). Bacteria belonged to 19 different phyla, with the most abundant being Bacteroidota (n = 2846), Firmicutes_A (n = 2696), Proteobacteria (n = 985), Firmicutes (n = 970) and Spirochaetota (n = 842) (**Figure 5A, 6A**).

**Figure 5:**
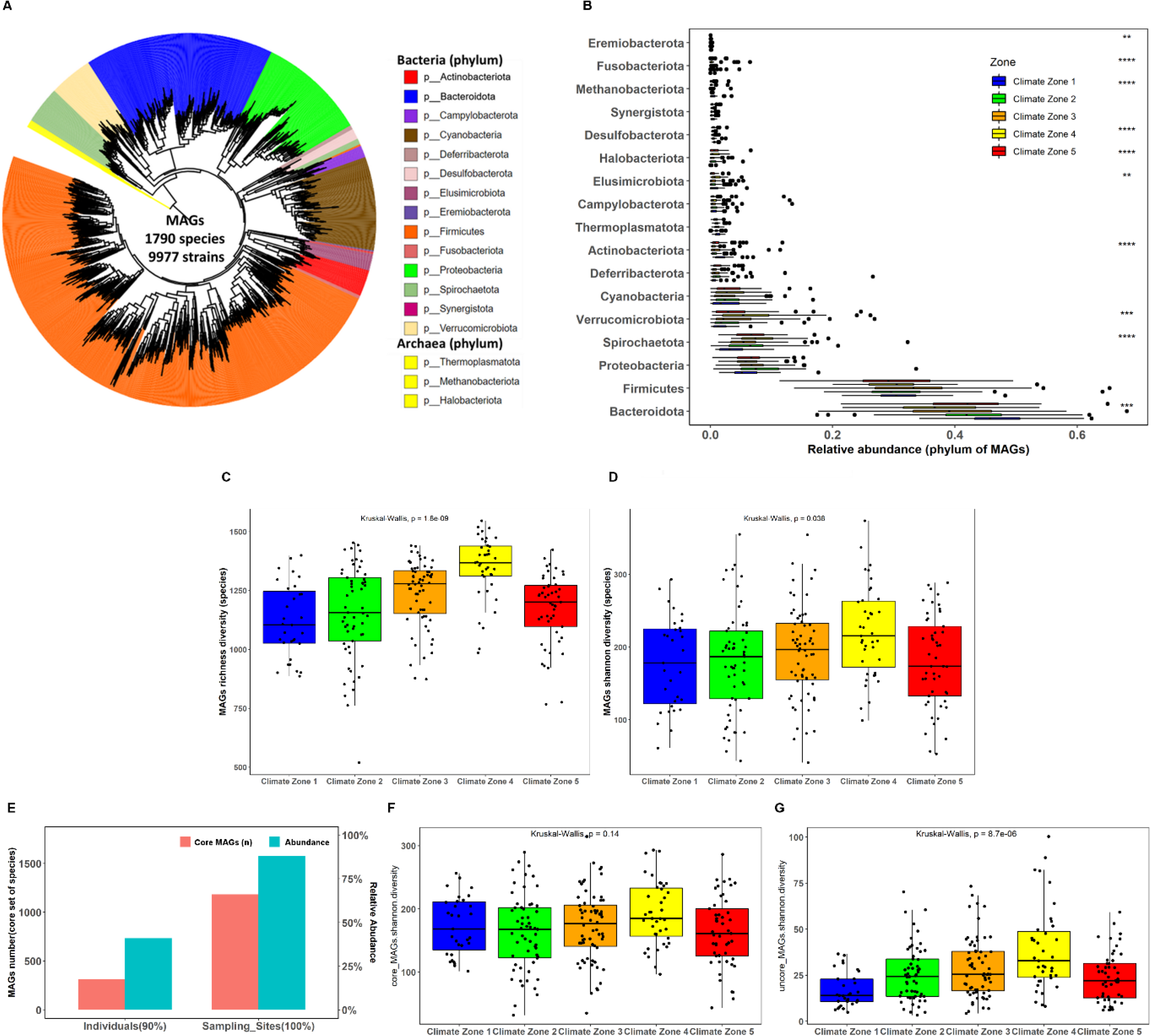
Species-level MAG taxonomy and diversity between samples and climate zones. To aid readability, all phyla classed as Firmicutes (Firmicutes_A, Firmicutes_B etc.) have been concatenated under the label “Firmicutes”. A) Phylogenetic tree of species-level MAGs. B) Abundance of phyla between climate zones (** p < 0.01; *** p < 0.001). C) Richness of MAGs between climate zones. D) Shannon diversity of MAGs between climate zones. E) The prevalence and relative abundance of core MAGs shared between 100% of sampling sites and at least 90% of individual samples. F) Shannon diversity of core MAGs between climate zones. G) Shannon diversity of non-core MAGs between climate zones.

Various bacteria of interest as food-borne pathogens or poultry pathogens were identified. *Campylobacter* spp. are a common cause of food-borne diarrhoeal disease in humans. Several species of *Campylobacter* were identified amongst the MAGs, including *Campylobacter avium* (15 strains), *Campylobacter coli* (4 strains), *Campylobacter jejuni* (10 strains) and two novel species. Various strains of *Escherichia* and *Shigella* are able to cause disease in humans through contaminated poultry meat. Surprisingly, only one strain of *E. coli* was identified among the MAGs, with most *Escherichia* strains identified as *Escherichia flexneri* (n = 17, also known as *Shigella flexneri*), and one strain being identified as *Escherichia dysenteriae*. Members of the *Helicobacter* genus are common causes of gastroenteritis. Ninety-five strains of *Helicobacter* were identified amongst the MAGs; 40 of these MAGs were identified as *Helicobacter pullorum*, with the remaining MAGs defined as belonging to 5 separate “*Helicobacter*” genera. Only one MAG was identified as belonging to the Chlamydia: *Chlamydophila gallinacean*.

In comparison to using the gene catalogue, after mapping raw sequencing reads to MAGs the proportion of reads annotated as bacterial and archaeal taxa were increased from 41.1% to 69.9% and from 0.4% to 1.3%, respectively (**Figure S4**). The relative abundance of MAGs in the samples was estimated, and the phylum abundance profiles were highly correlated between mapping using the gene catalogue or MAG database (r = 0.98). Using species-level MAG abundances, the diversity and richness of the caecal microbiota of chickens from different climate zones was compared. Around 66% of species were identified as core species, i.e. present in all 26 sampling sites. Far fewer core species were found when comparing individual samples (present in at least 90% of samples), but despite their smaller numbers these core species still accounted for around 50% of the average microbiota composition of samples (**Figure 5E**). The richness (**Figure 5C**, p = 1.8e-09) and diversity (**Figure 5D**, p = 0.038) of the MAGs differed significantly between climate zones. No significant differences were observed in alpha-diversity of the core species between climate zones (**Figure 5F**). However, for non-core members of the microbiota there were clear differences in alpha-diversity between climate zones (p = 8.7E-6) with diversity tending to decrease with altitude, except in climate zone 5 (**Figure 5G**).

We next wanted to identify whether any of our MAGs were differentially abundant in the three enterotypes that we had previously defined using our gene catalogue data. Of our 1790 species-level MAGs, 1404 were found to be differentially abundant between enterotypes (Kruskal Wallis adj-p < 0.05, **Supplementary table 3**). In total, one hundred and fourteen of these MAGs were found at 10-fold higher abundance in enterotype 1 in comparison to the other enterotypes. In contrast, only eight MAGs were found at 10-fold higher abundance in enterotype 2. Thirty-six significantly differently abundant MAGs were found at 10-fold higher abundance in enterotype 3 in comparison to the other enterotypes; these MAGs originated from a wide diversity of taxa (8 phyla, **Table 1**). We had previously identified *Prevotella* as being highly variable in abundance between climate zones and enterotypes. Of the fourteen *Prevotella* species that were differentially abundant between enterotypes, twelve were more abundant in enterotype 3 than in the other two enterotypes, with two MAGs at 10-fold higher abundance (*Prevotella sp000431975* and *Prevotella copri*).

**Table 1:** Species level MAGs that differed significantly in abundance between enterotypes

| Phylum | Total differentially abundant MAGs (n)* | Enterotype 1: 10-fold higher abundance (n)^ | Enterotype 2: 10-fold higher abundance (n)^ | Enterotype 3: 10-fold higher abundance (n)^ |
| --- | --- | --- | --- | --- |
| **Actinobacteriota** | **35** | **1** | **0** | **4** |
| **Bacteroidota** | **218** | **0** | **1** | **4** |
| **Campylobacterota** | **4** | **0** | **0** | **0** |
| **Cyanobacteria** | **129** | **19** | **0** | **3** |
| Deferribacterota | 5 | 0 | 0 | 0 |
| Desulfobacterota | 14 | 0 | 1 | 0 |
| Elusimicrobiota | 23 | 4 | 1 | 0 |
| Eremiobacterota | 2 | 1 | 0 | 0 |
| Firmicutes | 180 | 16 | 1 | 6 |
| Firmicutes_A | 510 | 48 | 3 | 11 |
| Firmicutes_B | 1 | 0 | 0 | 0 |
| Firmicutes_C | 14 | 0 | 0 | 4 |
| Firmicutes_G | 1 | 0 | 0 | 0 |
| Halobacteriota | 1 | 0 | 0 | 0 |
| Methanobacteriota | 1 | 0 | 0 | 0 |
| Proteobacteria | 147 | 21 | 1 | 3 |
| Spirochaetota | 52 | 2 | 0 | 1 |
| Synergistota | 1 | 0 | 0 | 0 |
| Thermoplasmatota | 9 | 1 | 0 | 0 |
| Verrucomicrobiota | 56 | 1 | 0 | 0 |
| Verrucomicrobiota_A | 1 | 0 | 0 | 0 |

*\*adj-p < 0.05, Species-level MAGs*
*^Relative abundance in comparison to other enterotypes. Only includes MAGs that were significantly differentially abundant*

### Comparing Ethiopian chicken MAGs to taxonomies found in non-scavenging chickens

The microbes identified in this study originate from scavenging chickens that are very distinct from commercial breeds, in terms of genetics, diets and environments. As such it may be expected that many of the microbes we have identified would be taxonomically and functionally distinct from those found in non-scavenging chickens. We therefore compared our MAGs to microbial genomes originating from a dataset of non-scavenging chickens (NSCs) and to the Genome Taxonomy Database (GTDB).

The majority of our strain-level and species-level MAGs were not present in either the NSC dataset or GTDB (**Figures 6B-C and 6E-F**, **Supplementary table 4**). For the strain-level MAGs, only 268 were identified in the NSC dataset and 47 in the GTDB, leaving 9682 strains that were not identified in either. At species-level, 423 of our MAGs were identified in the NSC dataset and 291 were identified in the GTDB, leaving 1242 species that were not identified. We clustered MAGs into 373 genus-level clusters, according to their amino acid identities (AAI). Of these genera, 163 were found in the NSC dataset while 266 were found in the GTDB, leaving 84 genera that were not identified in either (**Figures 6D and 6G**). Genera that were found to be unique to our dataset originated from a wide variety of taxa.

**Figure 6:**
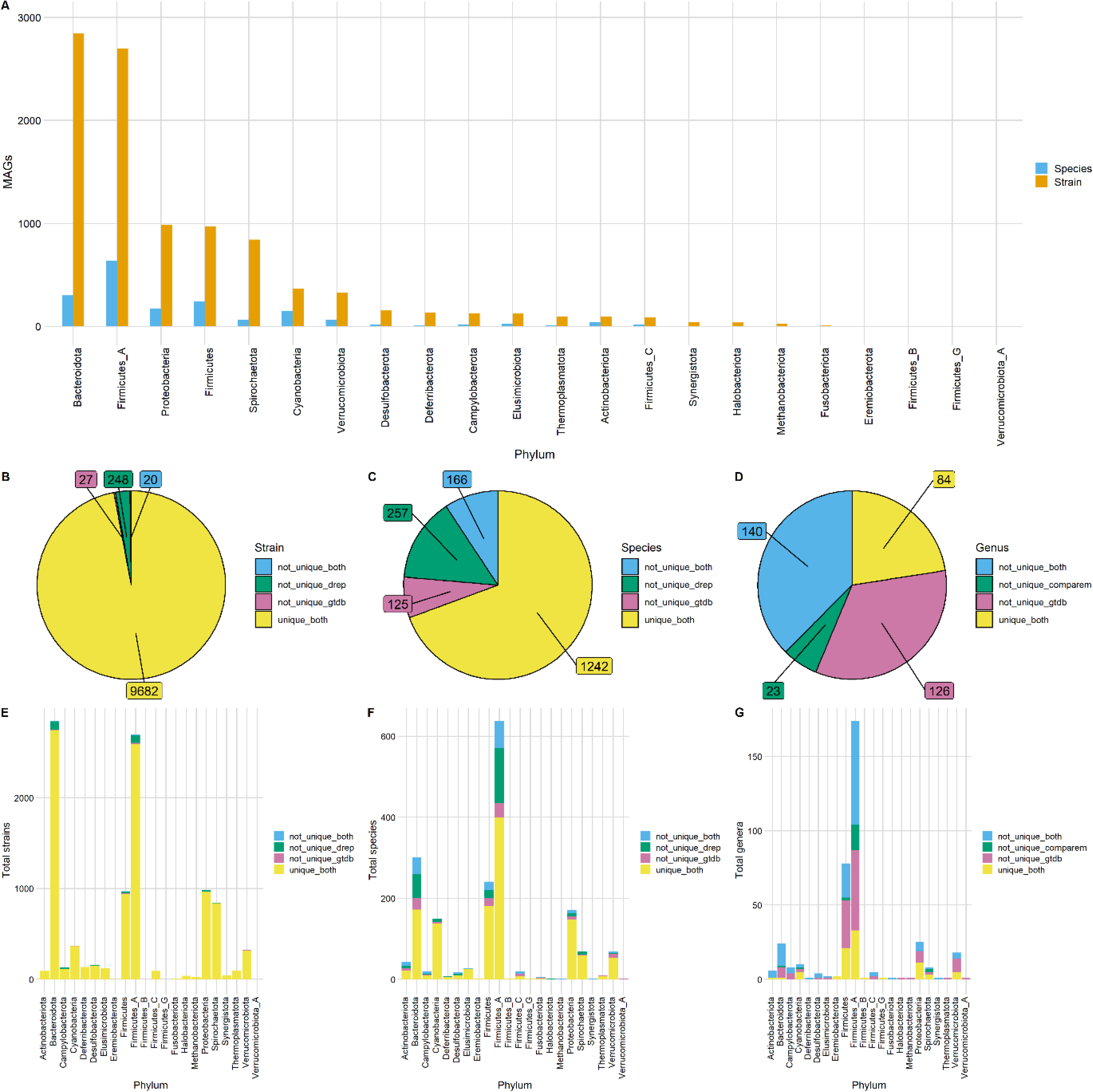
Comparison of Ethiopian chicken caecal MAGs to microbial genomes from non-scavenging chickens and the GTDB. A) Barplot showing the number of MAGS assigned to different phyla at both strain and species level. The number of strain-level MAGs (B and E), species-level MAGs (C and F) and genus-level clusters (D and G) that were assigned as “unique” to our dataset according to the following criteria: Strain and species level MAGs were not unique based on GTDB (“not_unique_gtdb”) if the average nucleotide identity (ANI) between the query and GTDB reference genome were > 99% or > 95% respectively. Genus-level clusters were not unique based on GTDB if any MAGs within that cluster were assigned taxonomy at genus level. MAGs were defined as not unique when compared to previous chicken microbial datasets (“not_unique_drep”) if they clustered at 99% (strain) or (95%) ANI with any NSC microbial genome. Genera were defined as not unique when compared to previous chicken microbial datasets (not_unique_comparem) if they clustered at 60% AAI with any NSC microbial genome.

The majority of species level MAGs carried at least one antimicrobial resistance (AMR) gene (n = 783), with tetracycline being the most commonly targeted drug class. Tetracycline resistance was also the most commonly targeted drug class among the NSC genomes. One MAG in particular, identified as *Escherichia flexneri*, was noted as containing a large number of AMR genes (n = 54) in comparison to other MAGs. Five of the NSC genomes also contained large numbers of AMR genes: *Pseudomonas aeruginosa* (n = 59), *Escherichia* sp. *Cla-CZ-1* (n = 56), *Escherichia whittamii* (n = 49), *Enterobacter roggenkampii* (n = 35) and *Klebsiella pneumonia* (n = 34).

### Functional characterisation of MAGs isolated from Ethiopian indigenous chickens

The caecal microbiota plays an important role in the fermentation of carbohydrates that are not able to be digested and absorbed in the small intestine of the host. Through this fermentation of fibrous compounds, SCFAs are produced. These SCFAs can be absorbed by the host and used as an energy source. It is likely that scavenging chickens consume different fibrous compounds and a greater diversity of fibre than non-scavenging chickens.

Our MAGs contained a large diversity of CAZymes (**Table S1**, **Supplementary table - MAGs metabolism strain (10.6084/m9.figshare.22154597) and Supplementary table - MAGs metabolism species (10.6084/m9.figshare.22154627)**). CAZymes are enzymes involved in the synthesis, metabolism and binding of carbohydrates. As such, it would be expected that a microbe that was rich in CAZymes would be able to thrive on a more diverse set of carbohydrates and would be more of a nutritional generalist than a microbe with a less rich CAZyme profile^18^. For total CAZyme composition: phyla clustered significantly by CAZyme composition for all domains (ANOVA: P = 1e-05, **Figure 7**).

**Figure 7:**
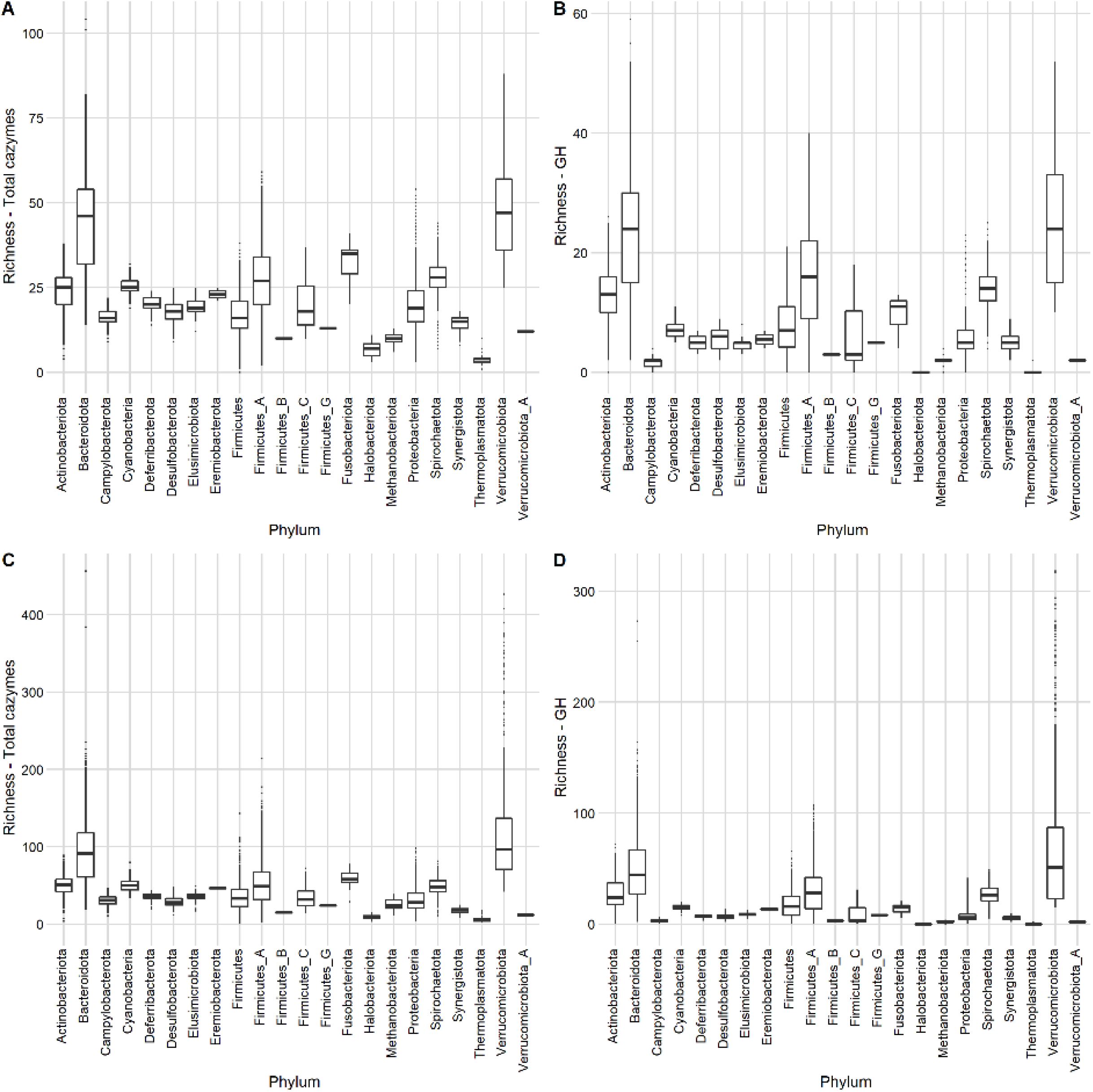
Boxplots showing number of CAZyme genes per strain-level MAG by phylum. A) Total unique CAZyme families. B) Total unique Glycoside Hydrolases (GH) families. C) Total CAZyme genes. D) Total GH genes.

Overall, MAGs from the Bacteroidota and Verrucomicrobiota contained the highest numbers of CAZyme genes, both by total CAZyme gene count (Bacteroidota: 92 ± 39, Verrucomicrobiota: 127 ± 84) and the number of unique CAZyme families (Bacteroidota: 44 ± 18, Verrucomicrobiota: 49 ± 15). For the Verrucomicrobiota, 41 MAGs contained over 250 CAZyme genes, and therefore represent some of the most CAZyme rich genomes in our dataset. These genomes belong to either of two families, Victivallaceae and UBA1829, and 16 were not identified in the GTDB or NSC dataset.

MAGs from the three Archaeal phyla had particularly low numbers of CAZymes: Thermoplasmatota (Total CAZyme genes: 6 ± 3, unique CAZyme families: 4 ± 1), Halobacteria (Total CAZyme genes: 9 ± 3, unique CAZyme families: 7 ± 2) and Methanobacteriota (Total CAZyme genes: 24 ± 8, unique CAZyme families: 10 ± 2). The MAGs with the lowest CAZyme richness (< 1 CAZyme genes) were members of the Mycoplasmatales (phylum: Firmicutes).

The average numbers of CAZymes per genome were similar between our dataset (60.08 ± 40.99) and the NSC dataset (68.24 ± 45.97) (**Figure S5**). In total, 266 CAZymes were shared between the two datasets, while nineteen were unique to the NSC dataset. A further nineteen CAZymes were identified in our MAGs but not in the NSC genomes: four carbohydrate binding molecules (CBM11, CBM65, CBM68 and CBM79), five glycoside hydrolases (GH47, GH86, GH107, GH119 and GH160), eight glycosyltransferases (GT13, GT15, GT40, GT44, GT60, GT74, GT75 and GT103) and two polysaccharide lyases (PL25 and PL32). These CAZyme genes originated from MAGs from a wide range of taxonomies. For example, for the glycoside hydrolases: GH47 (α-1,2-mannosidases) originated from two Bacteroidales strains, GH86 (β-agarase/β-porphyranase) originated from 8 strains of the family UBA1829 (phylum Verrucomicrobiota), GH107 (endo-α-1,4-L-fucanase) originated from one strain of the family UBA3636 (phylum Verrucomicrobiota), GH119 (α-amylase) originated from 2 Succinivibrionaceae strains, and GH160 originated from one Parabacteroides strain. Our MAGs also showed a wide diversity of predicted growth rates. These were found to significantly relate (P < 0.05) to CAZyme richness (**Supplementary Results 1**).

As well as identifying individual CAZyme genes present in MAGs, we can also use CAZyme data to identify which forms of carbohydrate are likely to be able to be digested by these strains by reconstructing metabolic pathways. Our MAGs demonstrated the capacity to degrade a wide variety of carbohydrates (**Figure 8, Figure S6** and **Supplementary table 5**). The most commonly encoded carbohydrate degradation pathway was for chitin, a polysaccharide commonly found in fungi and arthropods, (6828 of 9977 strain-level MAGs), which reflects the findings of the Distilled and Refined Annotation of Metabolism (DRAM) developers (48). This is closely followed by arabinose cleavage, present in 6648 MAGs.

**Figure 8:**
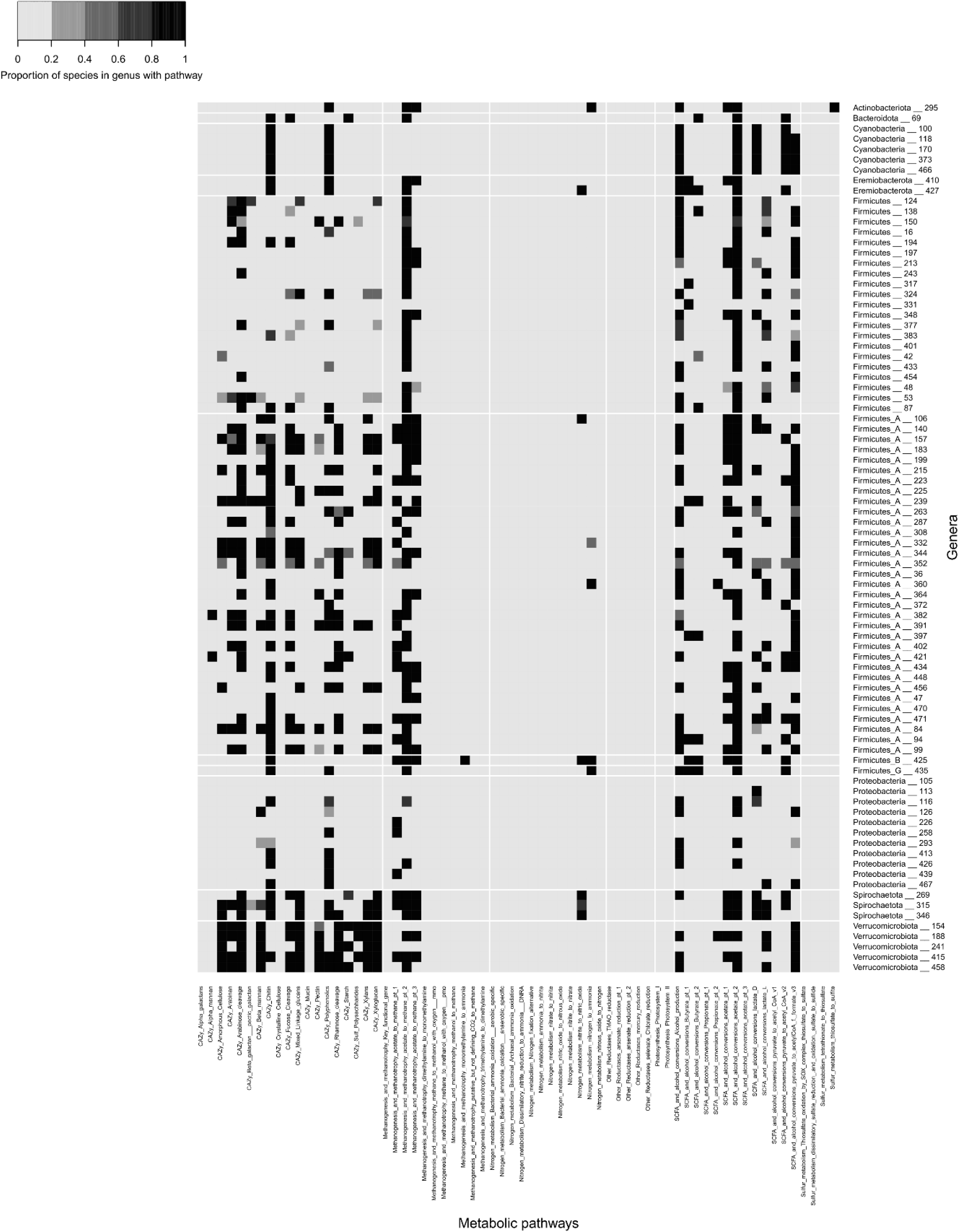
Heatmap showing the percentage of species-level MAGs within each ‘unique’ genera with particular metabolic pathways. Genera were clustered at 60% AAI using the output from CompareM. Genera were classified as ‘unique’ if no MAGs within that genus cluster were assigned a taxonomy at genus level by GTDB, and if no NSC genomes clustered at > 60% AAI with MAGs within that cluster.

Pathways to digest common fibrous plant compounds that are indigestible by chickens were present in many of the MAGs. Pathways for digestion of hemi-celluloses were very common, with mixed linkage glucan degradation encoded by 4689 MAGs, xyloglucans by 4420, xylans by 4151, beta-mannan by 3056 and alpha-mannan by 621. As expected from the diversity of CAZymes, members of the Bacteroidota were found to be the most likely to digest hemi-cellulose. In comparison to hemi-cellulose, the capacity to degrade amorphous cellulose (n = 3567) was less common. In general those phyla that contained more strains that could degrade amorphous cellulose were also more likely to contain strains that degraded hemi-cellulose and pectin. As well as plant/diet derived carbohydrates, microbes in the caeca have access to host derived carbohydrates such as mucin. Only 1.3% of strains showed the potential to degrade mucin.

Pathways for nitrogen metabolism were far less abundant amongst the MAGs than those for fibre degradation. MAGs from phyla that frequently harboured nitrogen metabolism pathways were rarely also able to degrade plant carbohydrates. For example, the majority of the members of the Desulfobacterota, Deferribacterota and Campylobacterota phyla harbour the dissimilatory nitrite reduction to ammonia (DNRA) pathway and/or were able to convert nitrate into nitrite, but less than 1% of these MAGs show any cellulose/hemi-cellulose degrading capacity. In contrast, members of the Spirochaetota were commonly able to metabolise nitrite to nitric oxide while also showing the capacity to degrade plant-derived carbohydrates.

While chickens produce little methane in comparison to other livestock, such as ruminants, members of their gut microbiota do have the capacity to carry out methanogenesis. Of the MAGs, 132 contained the key functional methanogenesis gene (methyl-coenzyme M reductase: mcr). Of those MAGs with the mcr gene, 22 were identified as having genes for all 8 steps required for methanogenesis, with a further 25 having at least 4 of the required steps. Interestingly, none of the genomes from the NSC dataset had genes for all eight steps required for methanogenesis, and only five genomes had > 50% of the required genes.

Gut microbes produce short chain fatty acids, principally butyrate, acetate, and propionate, by the fermentation of indigestible polysaccharides. These SCFAs can then be used as an energy source by the host animal. While it is difficult to be certain using metagenomic data whether a particular strain produces SCFAs, we can predict the potential for SCFA production using DRAM. The potential to produce SCFAs was widely encoded across taxonomies (**Supplementary table 5**). We visualised which SCFA/lactate encoding potentials occurred most together within the MAGs (**Figure 9**). By far the most common was the sole production of acetate, followed by the coproduction of acetate and lactate, then the coproduction of acetate and butyrate.

**Figure 9.**
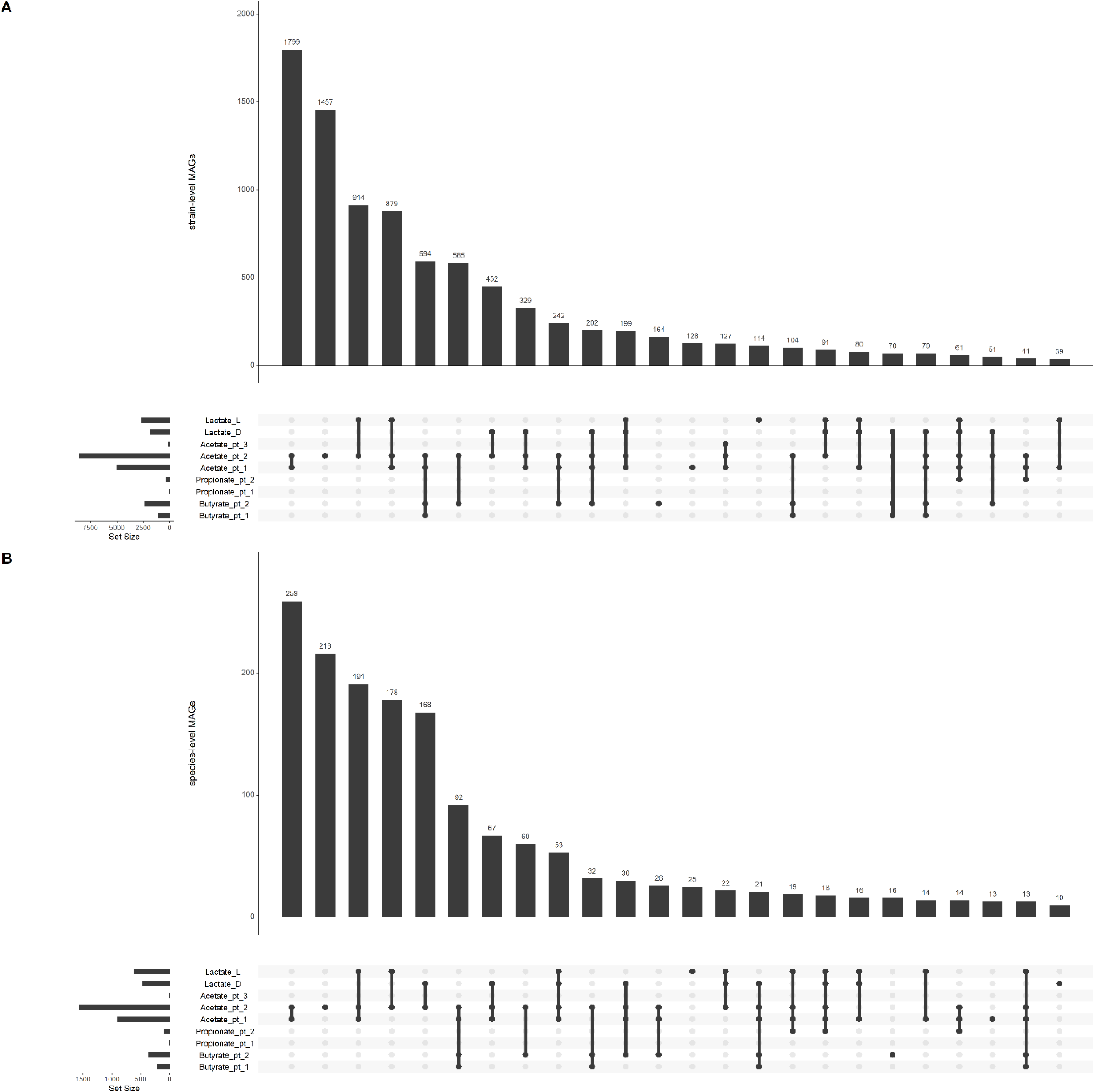
UpSet plots showing the number of MAGs with production potential for SCFA and lactate production, as defined by DRAM. Only includes intersections with ≥ 10 MAGs. A) Strain-level MAGs. B) Species-level MAGs.

## Discussion

In this study, we examined the caecal microbiota of 240 indigenous Ethiopian scavenging chickens originating from farms exposed to diverse climatic and geographic conditions. We constructed a gene catalogue containing 33 million genes, and 9,977 high-quality, strain-level MAGs originating from diverse taxonomies and with diverse functional capacities. We found that Ethiopian chicken caecal microbiota clustered into three distinct enterotypes, with one of these enterotypes being characterised by a high abundance of *Prevotella* and being particularly abundant in chicken living at high altitudes.

The chicken caecal microbiota is commonly dominated by bacteria, with low proportions of archaea and eukaryotes^19, 20^, which was reflected in our findings. Bacteroidota was the most abundant bacterial phylum in our samples, followed by Firmicutes, Proteobacteria and Spirochaetota. This reflects what has previously been found in scavenging/feral birds and adult hens, but contrasts with taxa commonly found in intensively-raised commercial broilers and young birds raised in biosecure poultry facilities. Commercial, intensively-raised chicken breeds are reared without contact with older chickens in highly biosecure poultry facilities.

They are therefore exposed to a lower diversity of microbial species than if they were raised by a maternal hen, or exposed to an outdoor environment. As such, these birds usually have a low-diversity gut microbiota dominated by Firmicutes until around two months of age, before developing a microbiota composition more similar to our findings by 30-50 weeks of age^21, 22^. Commercial broilers are commonly slaughtered at 5-6 weeks of age, and therefore do not have time to develop a “mature” microbiota. This is frequently also true for chickens that are part of microbiota studies, which often include only young birds. As such, the microbiota of these birds usually does not develop beyond a Firmicutes dominated composition^14, 23, 24^. This is in contrast to chicks raised by a hen or exposed to adult caecal/faecal contents, which have caecal microbiota more similar to our findings within a few days of hatch^25^.

At an inter-country level, geography has been demonstrated to have a significant impact on the chicken caecal microbiota^6^. We found significant differences in the alpha-diversity of the caecal microbiota between climate zones, with bacterial diversity generally decreasing as altitude increased. Our samples were also found to cluster into three enterotypes. Enterotypes are generally defined as a stratification of the gut microbiota based on similarities in terms of taxonomic compositions between samples^26^. Interestingly, one of our three enterotypes was predominantly associated with the climate zone with the highest altitude, and was characterised by a high prevalence of *Prevotella*.

Decreases in gut microbiota diversity have also been observed in a study of Tibetan chickens raised at different altitudes^8^. Tibetan chicken from high-altitudes also had a greater abundance of Prevotellaceae in their caeca, which correlates with our finding that *Prevotella* increases in abundance as altitude increases. This increase in *Prevotella* at high altitude has also been noted in house mice^27^, and cranes adapted to high altitudes have been demonstrated to have lower gut microbiota diversity than cranes that normally reside at low altitudes^28^.

Gut microbiota samples from pigs and humans living at high altitudes were also found to be significantly lower in diversity than those from low altitudes^29^. In contrast, the diversity of the gut microbiota of rhesus macaques was found to be increased in high-altitudes, and members of the Prevotellaceae were found in lower abundance^30^. The microbiota of humans living in high altitude areas of China have also been associated with increased microbiota diversity^31^; however, as in our samples *Prevotella* was found to be increased at higher altitudes. In chickens, while the reasons for this low diversity microbiota at high altitudes is currently unknown, it may be related to the effects of hypoxia^32^ or the relative lack of diversity in feed present at these elevations.

We compared our MAGs to a dataset made from microbial genomes previously constructed from the chicken gastrointestinal tract^13, 16, 33, 34^ (NSC dataset), and to the Genome Taxonomy Database^35^. The number and diversity of taxa that were identified in our dataset but not in the GTDB or NSC dataset emphasises the importance of studying the microbiota of indigenous livestock as well as commercial breeds. Our results also demonstrate the need to include microbial genomes isolated from indigenous livestock in commonly used genetic databases such as GTDB, as many of our MAGs were unable to be identified by GTDB-tk at the level of taxonomic order (n = 39), family (n = 204), and genus (n = 2252).

Despite the taxonomic novelty observed in our dataset, functional novelty was less obviously apparent. While there were significant differences between the types of CAZymes found in the NSC dataset vs our MAGs, the vast majority of CAZymes were found in both. This correlates with previous studies that found that differences in taxonomy tend to be greater than changes in function of the microbiota^36^. Particularly high numbers of CAZymes were found in the phyla Bacteroidota and Verrucomicrobiota, these phyla having been previously shown to have a high percentage of their total proteins being identified as CAZymes in comparison to other taxa^17, 37^. Several CAZymes were found in our MAGs, but not in the NSC dataset. These included several enzymes linked to the breakdown of algae: PL25 (ulvan lyase: 3 *Sphaerochaetaceae* MAGs), PL32 (poly(β-mannuronate) lyase / M-specific alginate lyase: 2 *Paludibacteraceae* MAGs), GH86 (β-agarase/β-porphyranase: 8 *UBA1829* MAGs) and GH107 (endo-α-1,4-L-fucanase: 1 *UBA3636* MAG)^38^. It is possible that this is due to the chickens consuming algae from local stagnant water sources.

As well as examining specific carbohydrate degrading enzymes, we also characterised the metabolic pathways present in our MAGs. The caeca is the main site of fibre fermentation in the chicken gastrointestinal tract. Both our MAGs and genomes from the NSC dataset contained pathways for the fermentation of a wide variety of fibrous compounds, including various forms of hemi-cellulose and amorphous cellulose. However, the ability to degrade crystalline cellulose was rare: encoded by only 6 MAGs. The result of fibre fermentation is the production of SCFAs, which can used by the host as an energy source, and can also play a role in pathogen resistance^10, 11^.

The SCFAs that had the most potential for production amongst our MAGs were acetate, then butyrate, then propionate, similar to the human gut^39^ and the NSC dataset. Several MAGs also showed the capacity to degrade mucin. Mucin degradation by gut microbiota can be related to a lack of dietary fibre, leading bacteria to rely on host-derived glycans^40^. However, some mucin-degrading species, such as *Akkermansia muciniphila* in humans, are often also common members of a healthy microbiota^41^. The caecal microbiota also plays an important role in nitrogen nutrition of the chicken, particularly when the bird is consuming little protein^42^, due to the reflux of urine into the caeca^11^. The capacity for nitrogen metabolism was present amongst our MAGs but far less so than plant fibre fermentation.

We have identified a wide functional and taxonomic diversity of microbes originating in the caecal contents of Ethiopian scavenging chickens under smallholder settings. The vast majority of these microbes were not identified in either the GTDB or a dataset of publically available microbial genomes isolated from predominantly non-scavenging chickens. We detected differences in the alpha and beta diversity of the chicken caecal microbiota relating to various climate and geographical factors. These findings highlight the potential hidden microbial diversity amongst indigenous, scavenging livestock that may be missed when examining only commercially reared animals.

## Methods

### Ethics approval and consent to participate

This study was reviewed and approved by the Institutional Animal Care and Use Committee (IACUC), International Livestock Research Institute (ILRI) (Reference number: ILRI-IREC2015-08/1).

### Sample collection

Two hundred forty three indigenous chickens were included in this study. They originated from 26 sites across 15 districts of Ethiopia that represent diverse agro-climatic conditions (**Figure 1A and 1B, Supplementary table 6 and Supplementary table 7**). Samples were collected from smallholder farms within 3 km^2^ of the sampling site centre (within 0.03 degrees of longitude or latitude). In order to avoid bias associated with household, each bird collected from a single site belonged to a different smallholder. Whole genome sequences had previously been produced for the majority of chickens in our study except for samples from Arginjona and Dehina_Maria that were mapped following previously described methods to the chicken reference genome GRRCg6a^43^. Principle coordinate analysis was conducted using PLINK v1.9^44^ to show clustering of chickens by autosomal single nucleotide polymorphisms (SNPs). SNPs were pruned for LD in PLINK (using PLINK option –indep-pairwise 50 10 0.1), then principle component analysis (PCA) plots were constructed in R. The chickens in our study did not belong to specific breeds; indigenous Ethiopian chickens do not constitute distinct breeds, instead weak sub-structuring of populations is observed based on geographic closeness^1^ (**Figure S7**).

Climate predictors were identified from the DIVA-GIS database based on a single geographic coordinate taken from the sampling site by GPS, as previously described^1^. Chickens were raised in a typical low input system, acquiring most of their food from scavenging and occasional supplementary feeding. Data was collected on rearing conditions, supplementary feeding, and local climate (**Supplementary table 7**). Most chickens lived in simple poultry houses, often constructed with a stone wall and grass roof, and with limited cleaning frequency (**Figure 1A**). All farmers practiced supplementary feeding, which included items such as kitchen waste, grains or vegetables. Chickens acquired most of their nutrition through scavenging on food sources located near to the household, including vegetation, insects, worms, wasted grains and animal faeces. Chickens frequently co-habited with other domestic animal species, and are also likely to have come into contact with wild animals.

### Clustering of climate zones

Climate predictions were obtained from WorldClim 2.0 Beta version 1 (June 2016). Clustering of climate zones was produced in R using the Kmeans cluster method based on annual temperature, annual precipitation and precipitation of the driest quarter of the sampling locations, during the years 1970 to 2000 (**Figure 1C and Figure S8**). This resulted in the clustering of sampling areas into five climate zones. A similar method for climate regionalization has previously been used by Yang *et al.*^45^.

### DNA extraction and shotgun metagenomic sequencing

Caecal contents were collected at the farm from scavenging chickens, and stored in RNAlater solution (Ambio). Samples were kept on ice for a maximum of 24 hours before being stored at −80°C prior to DNA extraction. DNA was extracted using the QIAamp Fast DNA Stool Mini Kit (Qiagen) following the manufacturer’s instructions, with some adjustments as described previously^46^, including the addition of a bead beating step and increasing the cell lysis temperature from 70°C to 95°C to increase the likelihood of lysing Gram-positive bacteria. DNA sequencing libraries were prepared using the Nextera XT DNA Library Prep Kit (Illumina) and sequenced with an Illumina Novaseq (2×150 bp) (Berry Genomics Co.). All 243 samples were sequenced in a single run, yielding up to 10 Gb per sample. Adapter trimming and quality filtering were performed using Fastp^47^ (v.0.1.24). Host reads were removed by mapping the *Gallus gallus* genome (GRCg6a) to the trimmed sample fastq files using BWA-MEM (v.0.7.15)^48^, followed by SAMtools (v.1.3.1)^49^ to select reads where both paired-end reads were unmapped. Three samples contained high host contamination and were thereby not taken forward for further metagenomic analysis (except for construction of MAGs), leaving a total of 240 samples.

### Gene catalogue construction and analysis

After quality control and host removal, a gene catalogue was constructed from non-redundant genes. Firstly, single-sample assembly was conducted using MEGAHIT (v.1.2.9) (contig length > 500bp)^50^. Then, MetaGeneMark-1^51^ (GeneMark.hmm v.3.38: gmhmmp -a -d -f G -m; MetaGeneMark_v1.mod; gene length > 200nt) was used to predict open reading frames (ORFs). Non-redundant genes were clustered using CD-HIT^52^ (v.4.6.6) at 95% identity over 90% of the shorter ORF length (-c 0.95 -aS 0.9 -M 0 -T 0), resulting in 33,629,587 genes. As many as 97.9% (95.2∼98.5%) of the sequencing reads could be included in the non-redundant gene catalogue. Dual-BLAST least common ancestor strategies were used for microbiome taxonomic annotation. Diamond^53^ (v.2.0.2; diamond blastp -c 1 -k 5 -f 6) was used to search for homologous genes in the Uniprot database (version 2019_03) using the protein sequences in the non-redundant gene catalogues. Then, the aligned Uniprot regions (E-value < 10−5) were used in a second alignment against the Uniprot database, resulting in the identification of homolog neighbourhoods of the initial query genes by reporting e-values that were equal to or less than the e-value from the first alignment. Query genes were then assigned the taxonomy of the least common ancestor of the neighbourhood. Sequencing reads were directly mapped to the non-redundant gene catalogues by BWA-MEM (v.0.7.12 - default options)^48^. The relative abundance of genes in the samples was calculated based on the count of aligned reads, normalized to the gene length and sum of the abundance of all genes. Based on taxonomic annotation of the gene catalogue, aligned reads were used to quantify taxonomic abundance profiles within samples. Relative taxonomic abundance was normalized to sequencing depth and the length of the genes originating from the same level of taxonomic classification.

### Enterotype clustering

Gut enterotype analysis was performed by multidimensional cluster analysis and PCA on genus abundance, according to previously described methods^54^, using Jensen–Shannon divergence. All genera from the gene catalogue were used to define enterotypes in order to include taxa from all kingdoms. LEfSe was used to determine the microbiota features that most likely explained differences between enterotypes^55^. Spearman correlations were performed between the main genera contributing to enterotype clustering and all other genera. Genus networks were visualized using the Cytoscape^56^ platform by transforming Spearman correlations (p < 0.01, rho > 0.5) into links. Mantel tests (Spearman correlation) were performed between our previously identified non-redundant environmental factors and genera and phyla. Spearman pairwise correlations between continuous metadata variables were calculated (SparCC)^57^. P-values were corrected for multiple testing using Benjamini-Hochberg False Discovery Rate (FDR) correction. Significant features were used as input for building linear models using stepwise regression based on the Akaike Information Criterion^58^. The Shannon diversity index and inverse Simpsons index were used to compare alpha-diversity between groups (Kruskal-Wallis). Bray-Curtis dissimilarity values were calculated to assess beta-diversity, and PERMANOVA was used to compare groups.

### Metagenome assembled genome assembly

Metagenomic bins were constructed using two different methods:

Method 1: Coassemblies and single-sample assemblies were performed using MEGAHIT (v1.1.3) with a minimum contig length of 500 bp. Using MEGAHIT, five separate coassemblies were performed on samples from each climate zone. BWA MEM was used to map reads from each sample to the assembly from the same sample. MetaBAT2 (option -m 1500) was applied for contig binning.

Method 2: All 243 samples were used for sequence assemblies. IDBA-UD (v.1.1.3)^59^ was used for single sample assembly (options: --num_threads 16 --pre_correction --min_contig 300). BWA MEM was used to map reads from each sample to the assembly from the same sample. SAMtools was used to create BAM files, and coverage for each assembly was calculated by running the command jgi_summarize_bam_contig_depths on these files. MEGAHIT was used for coassembly of all samples (v.1.1.1) (options: --kmin-1pass -m 100e+10 --k-list 27,37,47,57,67,77,87 --min-contig-len 1000), in six randomised batches of samples. Contigs were filtered to a minimum length of 2kb, and mapped as for single assemblies. MetaBAT2 (v.2.11.1)^60^ was used to construct metagenomic bins for both single sample assemblies and coassemblies (options: --minContigLength 2000, --minContigDepth 2).

The completeness and contamination of bins were assessed using CheckM^61^ (v.1.1.3; options: lineage_wf -t 30 -x fa --nt --tab_table). Bins with completeness ≥ 80% and contamination ≤ 5% were concatenated and use as an input for DAS Tool^62^ (v1.1.2; option: --search_engine diamond -c merge.contigs.fa --threads 12 --write_bins 1), as an additional quality control step. The bins output by DAS tool were dereplicated using drep^63^ at 99% average nucleotide identity (ANI), which is equivalent to a microbial strain, and at 95% ANI, equivalent to a microbial species. Dereplication is the process of reducing a set of genomes based on their sequence similarity. GTDB-Tk^64^ (v.0.3.2, Database release 95) was used to assign taxonomy to MAGs. The taxonomic names of phyla used throughout the text of this manuscript are based on those used in this version of GTDB-Tk. Phylogenies were constructed using Phylophlan 3.0^65^ (v.3.0.60; options:-d phylophlan --min_num_markers 60 --subsample phylophlan -f tol_config.cfg --diversity high --fast --genome_extension fa –nproc 2). Phylogenetic trees were visualised using Graphlan^66^ (v.1.1.3.1) and iTOL^67^ (Interactive Tree Of Life; v.5). The table2itol package (https://github.com/mgoeker/table2itol) was used for phylogenetic tree annotations.

MAG genes were identified using Prodigal^68^ (v.2.6.3). CD-HIT^52^ (v.4.6.8) was used to cluster all MAG proteins at 100% similarity and 90% similarity. The relative abundance of MAGs in the samples was estimated using the quant_bins module of MetaWRAP (v.1.3) with default parameters^69^. Kruskal-Wallis with Benjamini-Hochberg FDR correction was used to identify species-level MAGs that were differentially abundant between enterotypes.

### Comparing MAGs to previous chicken microbiota datasets

We constructed a dataset of chicken derived microbial genomes from previously published chicken microbiota studies (**Supplementary table 8**): including 5,595 MAGs and 41 genomes of representative cultured isolates of novel species from Gilroy *et al.*^13^; 469 MAGs from Glendinning *et al.*^16^; 133 genomes of cultured isolates from Medvecky *et al.*^33^; and 16 genomes of cultured isolates from Zenner *et al.*^34^. These genomes were dereplicated using dRep (v.3.2.2) (options: -comp 80 -con 10 -str 100 -strW 0) at 99% ANI (strain-level) and 95% ANI (species-level) to produce two datasets of dereplicated, high-quality genomes. The vast majority of samples from which these genomes were isolated originated from non-scavenging chicken (except 22 (prior to dereplication) sampled from NCBI Bioproject PRJNA616250). We therefore labelled these datasets NSC (non-scavenging chickens) and compared our MAGs to these datasets to assess whether they were taxonomically and functionally distinct.

To calculate whether our MAGs were taxonomically distinct at strain and species level, dRep was used on our MAGs and the NSC datasets at 99% and 95% ANI. Any MAG which did not cluster at 99% ANI with any genome from the NSC dataset was classed as distinct at strain level. Any MAG which did not cluster at 95% ANI with any genome from the NSC dataset was classed as distinct at species level. Species-level genomes from both our dataset and the NSC dataset were compared using CompareM^70^ (v.0.1.2) aai_wf to generate average AAI. Genera were clustered at > 60% AAI; genera were defined as distinct if they contained no NSC genomes. Our MAGs were also compared to the GTDB^64^: this database is commonly used to taxonomically classify bacterial and archaeal genomes. MAGs were defined as distinct strains in comparison to genomes in the GTDB if the ANI output by GTDB-Tk was < 99%, and distinct species if the ANI output by GTDB-tk was < 95%. Genera clusters (> 60% AAI) were defined as distinct from those in the GTDB if no MAG within that cluster was assigned a genus by GTDB-Tk.

### Functional annotation of MAGs

Our MAGs and genomes from the NSC dataset were annotated in order to understand the potential metabolic function of these microbes. Genomes were annotated using DRAM^39^ (v.1.2.2) with the ‘annotate’ command; these annotations were then curated and summarised using the ‘distill’ command. DRAM is a tool for annotating MAGs, using various databases, including UniRef90^71^, PFAM^72^, dbCAN^73^, RefSeq viral^74^, VOGDB (http://vogdb.org/) and the MEROPS peptidase database^75^, and a user supplied version of the KEGG^76^ database (downloaded Sep 15th 2018). This tool provides information on the overall metabolic pathways encoded by the genomes, as well as specific information on the presence of CAZymes, rRNA genes and tRNA genes. Permutational multivariate analysis of variances (PERMANOVAs) were conducted using the adonis command from Vegan^77^ (v.2.5.7), to compare the types of CAZymes present between groups. The Kruskal-Wallis test was used to compare the abundance of CAZyme genes between groups. Microbial growth rates were predicted using gRodon^78^ (v.0.0.0.9). This tool estimates maximal microbial growth rates by comparing codon usage patterns in highly expressed genes versus other genes. We first used prokka^79^ (v.1.14.6) (options: --centre X –compliant) to identify genes, including highly expressed genes (genes annotated as ribosomal proteins). Predicted genes and highly expressed genes were used as input for the gRodon ‘predictGrowth’ command, which was ran in partial mode to account for incomplete genomes. As suggested by the gRodon creators^78^, copiotrophs were defined as having a < 5 hours doubling time, whereas oligotrophs were defined as having > 5 hours doubling time. AMR genes were identified in species level MAGs using the Resistance Gene Identifier (v. 5.1.1), and Comprehensive Antibiotic Resistance Database reference sequences downloaded on 30^th^ March 2021^80^.

### Selection of environmental variables

The relationships between environmental variables and the abundance of microbial taxa were estimated. In order to reduce model complexity by limiting the amount of environmental variables included in our analyses, variance inflation factors (VIF) were calculated for a set of environmental variables (**Supplementary table 9**), to identify collinearity among explanatory variables (VIF value less than 20). The highest contributing set of uncorrelated environmental variables was identified. This led to the selection of 24 variables that were included in our analyses: 8 climate/geographical variables (altitude, bio2, bio3, bio13, bio14, bio15, bio18, bio19), 10 soil/land cover factors (CULT, FOR, SNDPPT, SLTPPT, CRFVOL, BLDFIE, CECSOL, ORCDRC, PHIHOX, WATCAP) and the dominance of 5 feeding crops (Wheat, Maize, Barley, Millet, Teff) and Ingera (a food derivative from Teff). Redundancy analysis (RDA; Vegan 2.6.2 package) was used to identify environmental variables that contributed to microbiota variation.

### Graphical analyses

Graphs were created in R. Plots were constructed using the packages ggplot2^81^, Cowplot^82^, UpsetR^83^, cluster, clusterSim, ggpurb and corrplot. Heatmaps were constructed using heatmap.2 from gplots^84^ (v. 3.0.1).

## Data availability

The paired-read fastq files and species-level MAG fasta files generated and analysed during the current study are available in the European Nucleotide Archive under project PRJEB57055. Strain-level MAG fasta files (doi:10.6084/m9.figshare.22140284) and the gene catalogue (doi: 10.6084/m9.figshare.22180336) are available through figshare. Metabolic data on species level (doi: 10.6084/m9.figshare.22154627) and strain level (doi:10.6084/m9.figshare.22154597) MAGs are available through figshare.

## Supporting information

Supplementary Results 1

## Acknowledgements

We thank ILRI staff Michael Tesmegen for field support, and Mick Watson for advice on the analysis of our data.

## Author contributions

LG and XJ contributed equally to the data analysis and writing of the paper. OH, JH and XJ designed the study. AK, JH, OH, JBH, JEP, and WP organised and/or collected the samples in the field. SO contributed to study oversight and performed the DNA extraction of the samples. JBH, KK, HJ, and OH contributed to the writing of the paper. AA contributed to the data analysis. All authors read and approved the final manuscript.

