## Supplementary Results 1 for "Altitude-dependent agro-ecologies impact the microbiome diversity of scavenging indigenous chicken in Ethiopia"

**Supplementary Results 1: Detection of slow-growing microbes**

One advantage of metagenomics over culturomics is that culturing often leads to an overrepresentation of faster growing microbes. We wanted to assess the degree to which growth rate varied amongst our MAGs, and particularly whether we were able to detect slow growing strains that would be more likely to be missed by culturing alone. Maximal growth rate is frequently used to quantify the growth of microorganisms, referring to the doubling time (dt) under optimal nutritional conditions. It is possible to estimate the maximal growth rate of a microbe using genome-wide codon usage statistics^1^. Above a 5 hour minimal dt maximal growth rate tends to be underestimated as above this point less signal for selection of growth optimisation is observed. This has been proposed as a threshold point for defining copiotrophs (<5 hours dt) from oligotrophs (>5 hours dt)^1^. As the environment of the chicken caeca is abundant in nutrients it would perhaps be expected that the majority of the microbiota would be copiotrophic. Our MAGs displayed a wide range of maximal growth rates (**Figure 1**). As previously noted by Weissman *et al.*^1^, members of the Actinobacteria (1.28 ± 0.51 dt, 100% copiotrophs), Bacteroidota (3.1 ± 1.8, 85% coliotrophs), Firmicutes (Firmicutes _A: 3.1 ± 1.7 dt, 88% coliotrophs, Firmicutes: 2.8 ± 1.4 dt, 93% copiotrophs) and Proteobacteria (3.2 ± 2.2 dt, 75% copiotrophs) were predominantly faster growing. Unlike the Weissman *et al.* study, members of the cyanobacteria were found to be predominantly fast-growing (4.0 ± 1.8 dt, 82% copiotrophs). Members of the Camplyobacteriota were noted as having particularly high doubling times and being predominantly classified as oligotrophs (84.5% oligotrophs).

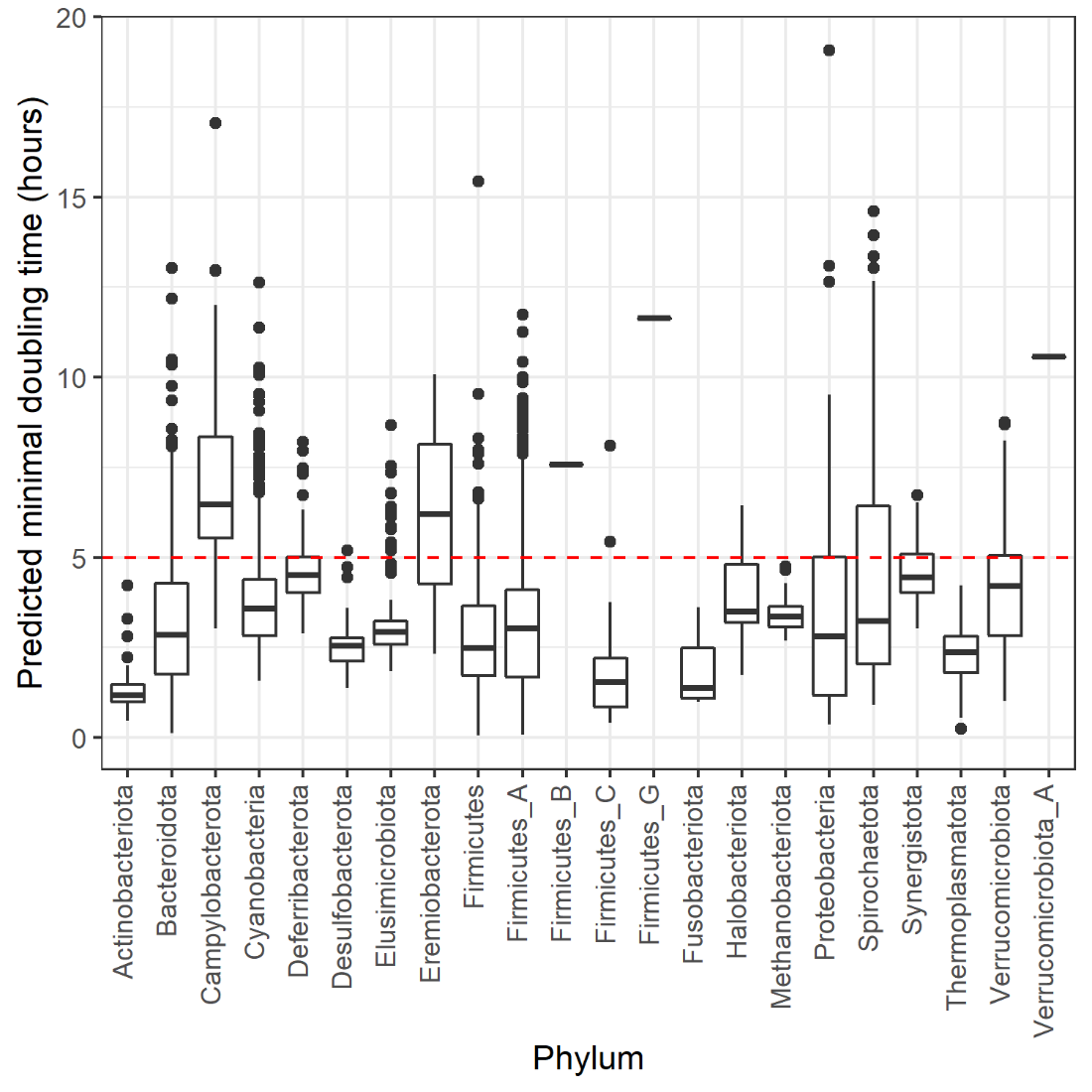

**Figure 1: Boxplot showing the predicted doubling time of strain-level MAGs, by phylum. The red, dashed line indicates the cut-off point above which MAGs are considered oligotrophs.**

We also compared the growth rate of our MAGs to the NSC dataset. In order to assess the potential impact of whole genome sequencing of cultured isolates vs metagenome assembled genome construction, we grouped winning genomes from the 99% ANI dereplication of the NSC dataset and separated them into those that originated from cultured isolates (NSC_cultured: 170 genomes) and those that were constructed from metagenomic data (NSC_MAGs: 2343 genomes). Cultured isolates contained a greater proportion of fast-growers than either the NSC_MAG group or our MAGs (**Figure 2).**

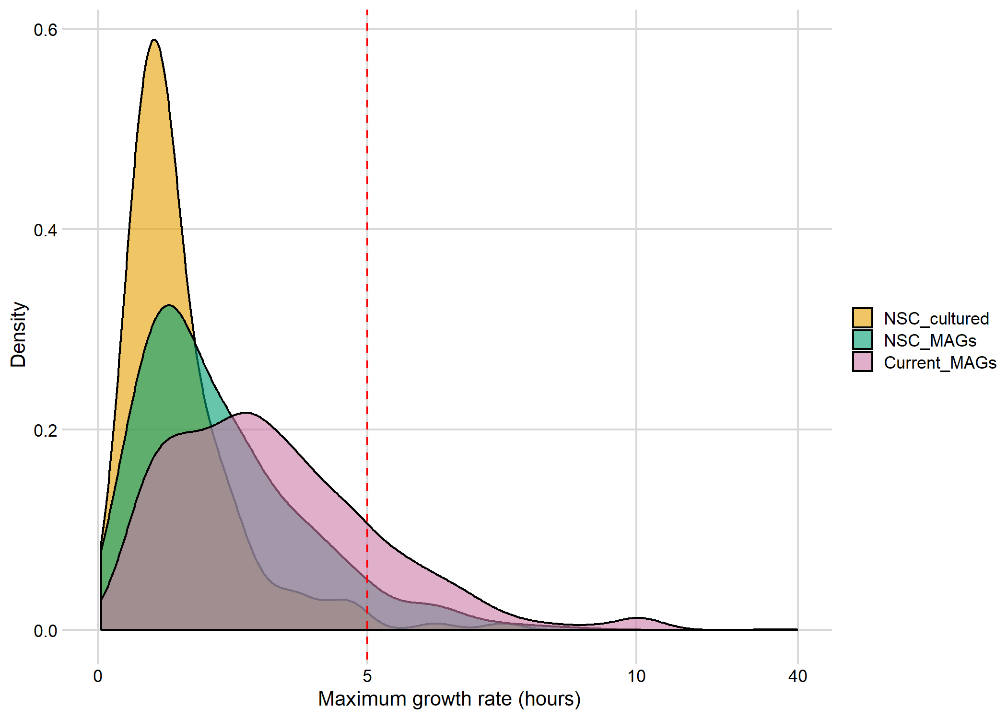

**Figure 2: Growth rate distributions for strain-level genomes from MAGs produced in this study (Current_MAGs: 9977) and dereplicated genomes from the NSC dataset, which originated from cultured isolates (NSC_cultured: 170) or were MAGs (NSC_MAGs: 2343). The red, dashed line indicates the cut-off point above which MAGs are considered oligotrophs.**

**Relating CAZyme abundance to oligotrophy**

We also wanted to test whether growth rate was related to CAZyme richness, as oligotrophy is generally related to substrate specialisation whereas copiotrophy is related to substrate generalisation. It would therefore be expected that specialists would have lower CAZyme diversity, and therefore growth rate would be inversely related to CAZyme richness. CAZyme composition significantly differed based on whether a MAG was defined as an oligotroph or coliotroph (ANOVA P<0.001) and significant differences (Kruskal-Wallis: P<2.2e-16) in CAZyme richness (total and unique CAZyme counts) were identified between oligotrophs (Total CAZyme genes: 50.8 ± 36.3, Unique CAZyme families: 26.4 ± 13.3) and copiotrophs (Total CAZyme genes: 62.0 ± 41.6, Unique CAZyme families: 31.1 ± 15.3) (**Figure 3**).

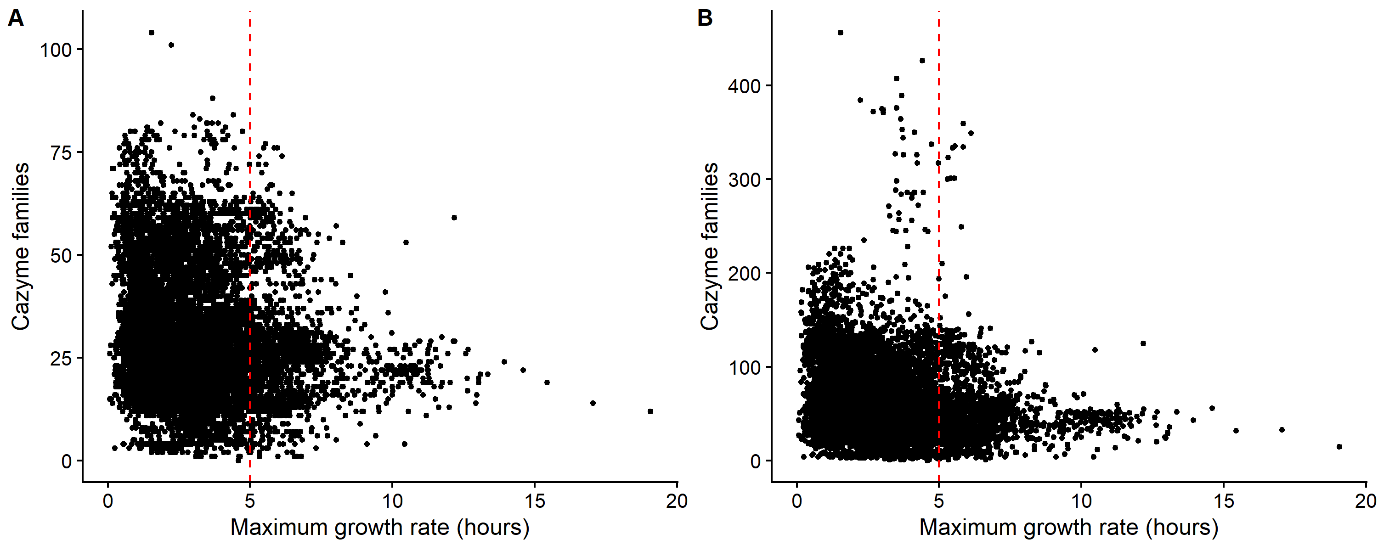

**Figure 3: Total CAZyme families count vs maximal growth rate. A) Unique CAZyme families, B) Total CAZyme genes. The red, dashed line indicates the cut-off point above which MAGs are considered oligotrophs. Copiotrophs were found to have significantly more CAZyme genes and unique CAZyme families than oligotrophs (Kruskal-Wallis: P<2.2e-16).**
