## Supplementary Figures and Tables for "Altitude-dependent agro-ecologies impact the microbiome diversity of scavenging indigenous chicken in Ethiopia"

**Figure S1:**

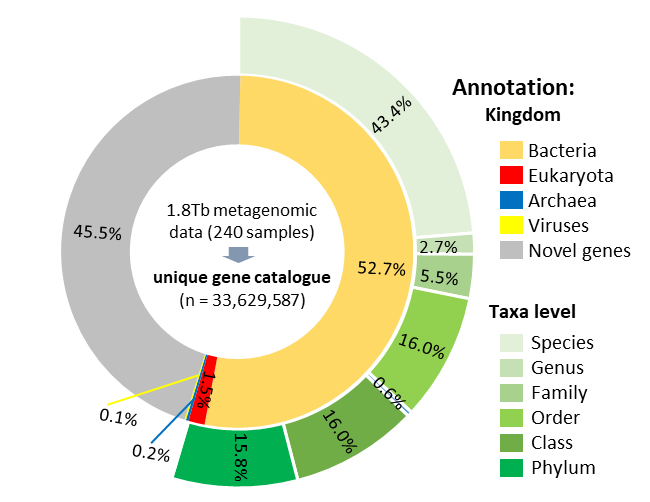

**Construction of a chicken caecal reference gene catalogue. The 3,629,587 non-redundant genes contained in the catalogue represent the metagenomes of 240 chicken caecal contents samples. Non-redundant genes were assigned to different taxon levels based on their last common ancestor in the Uniprot database (version 2019_03).**

**Figure S2:**

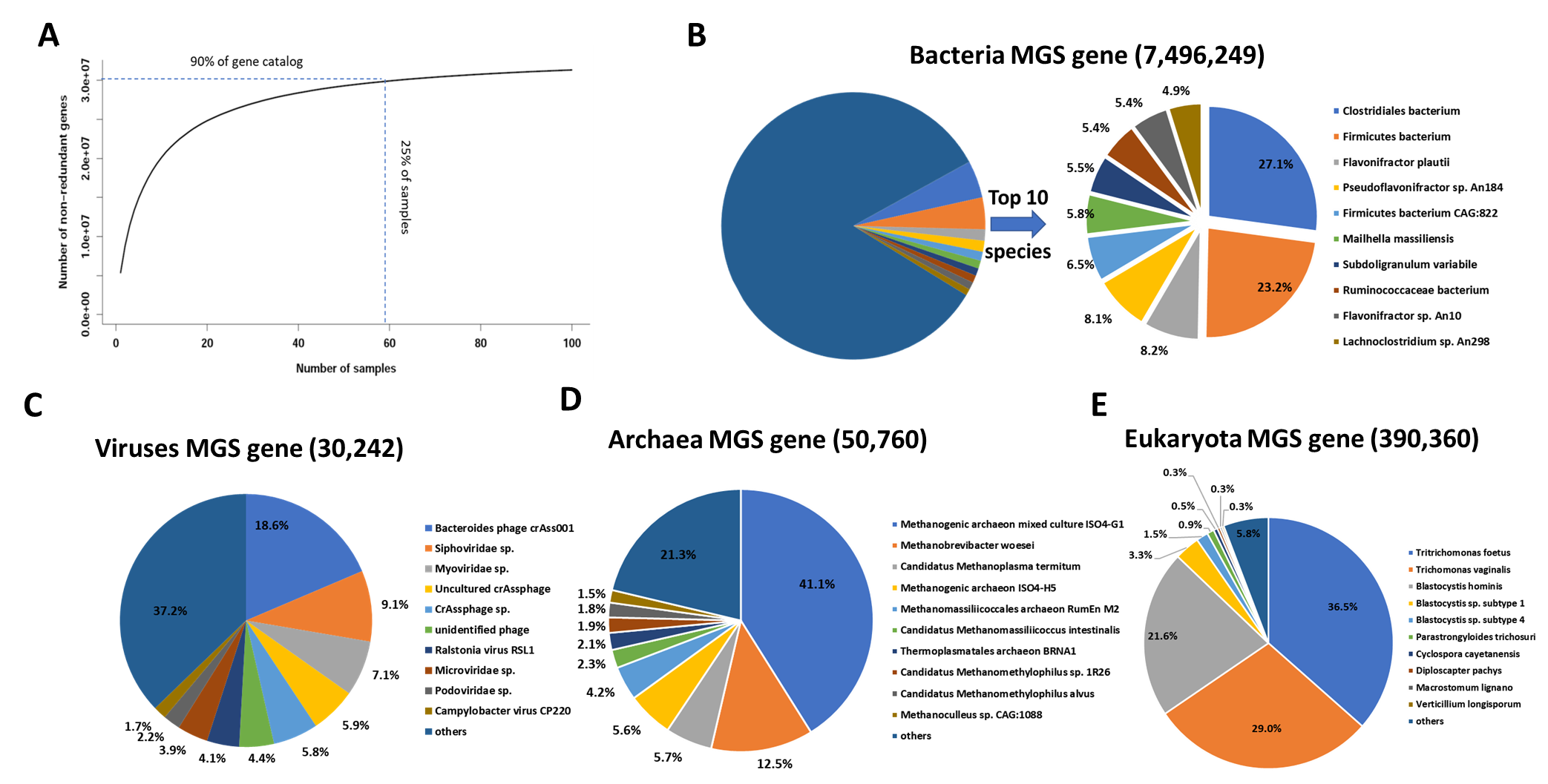

**Description of the gene catalogues constructed from the caecal microbiota of Ethiopian indigenous chickens. A) Rarefaction analysis of the number of non-redundant genes vs sampling number. B-E) Break down of the taxa identified in the Ethiopian chicken caecal microbial gene catalogue, by Kingdom.**

**Figure S3:**

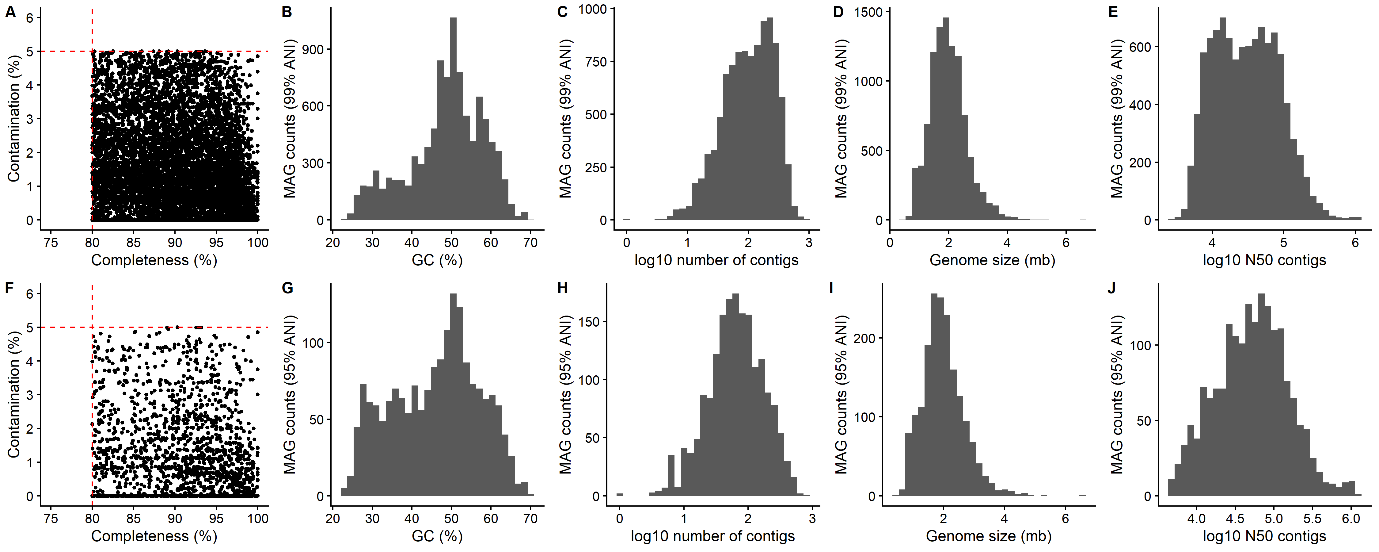

**Genome statistics of high-quality, non-redundant strain-level (A-E) and species-level (F-J) metagenome assembled genomes, as defined by CheckM. A and F: Completeness and contamination – dashed red lines indicate cutoffs for defining genomes as high-quality. B and G: Percentage GC content. C and H: log10 number of contigs per genome. D and I: Genome size (mb). E and J: log10 N50 of contigs.**

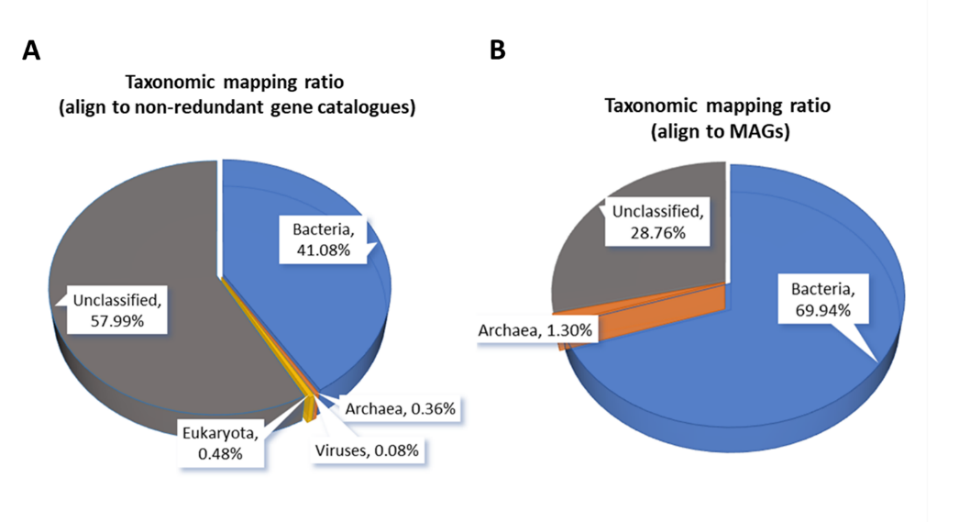
**Fig S4:**

**The proportions of anotated read after mapping raw sequencing reads to the non-redundant gene catalogue (A) and MAGs (B).**

**Figure S5:**

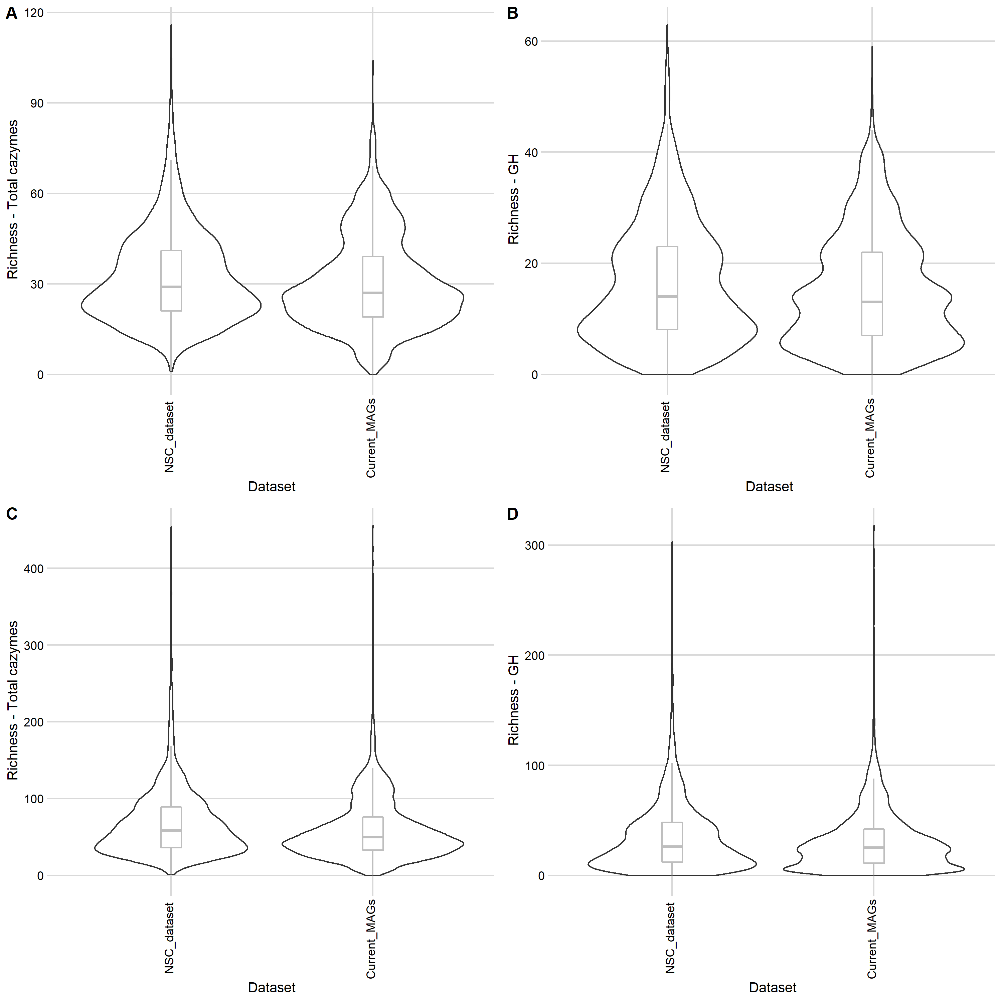

**Violin plot showing number of CAZyme genes per strain-level MAG by dataset. A) Total unique CAZyme families. B) Total unique Glycoside Hydrolases (GH) families. C) Total CAZyme genes. D) Total GH genes.**

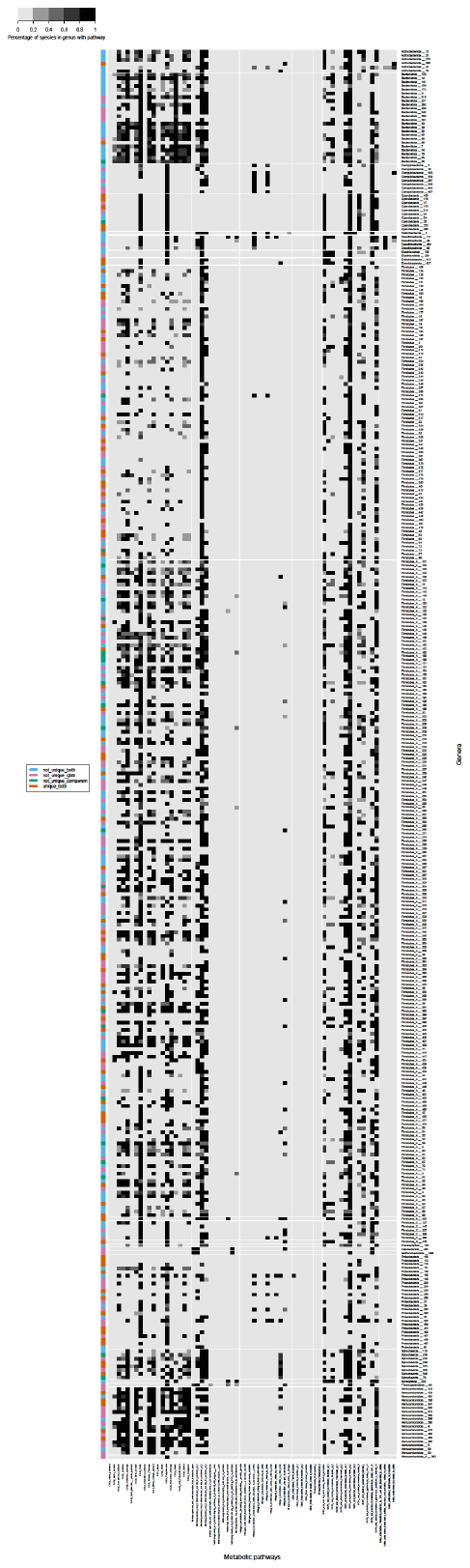

**Figure S6: Heatmap showing the percentage of species-level MAGs within genera with particular metabolic pathways. Genera were clustered at 40% AAI using the output from comparem. The uniqueness of genera in comparison to previous datasets are indicated. Genus-level clusters were not unique based on GTDB if any MAGs within that cluster were assigned a taxonomy at genus level. MAGs were defined as not unique when compared to previous chicken microbial datasets (“not_unique_drep”) if they clustered at 99% (strain) or (95%) ANI with any non-scavenging chickens (NSC) microbial genome. Genera were defined as not unique when compared to previous chicken microbial datasets (not_unique_comparem) if they clustered at 60% AAI with any NSC microbial genome.**

**Figure S7:**

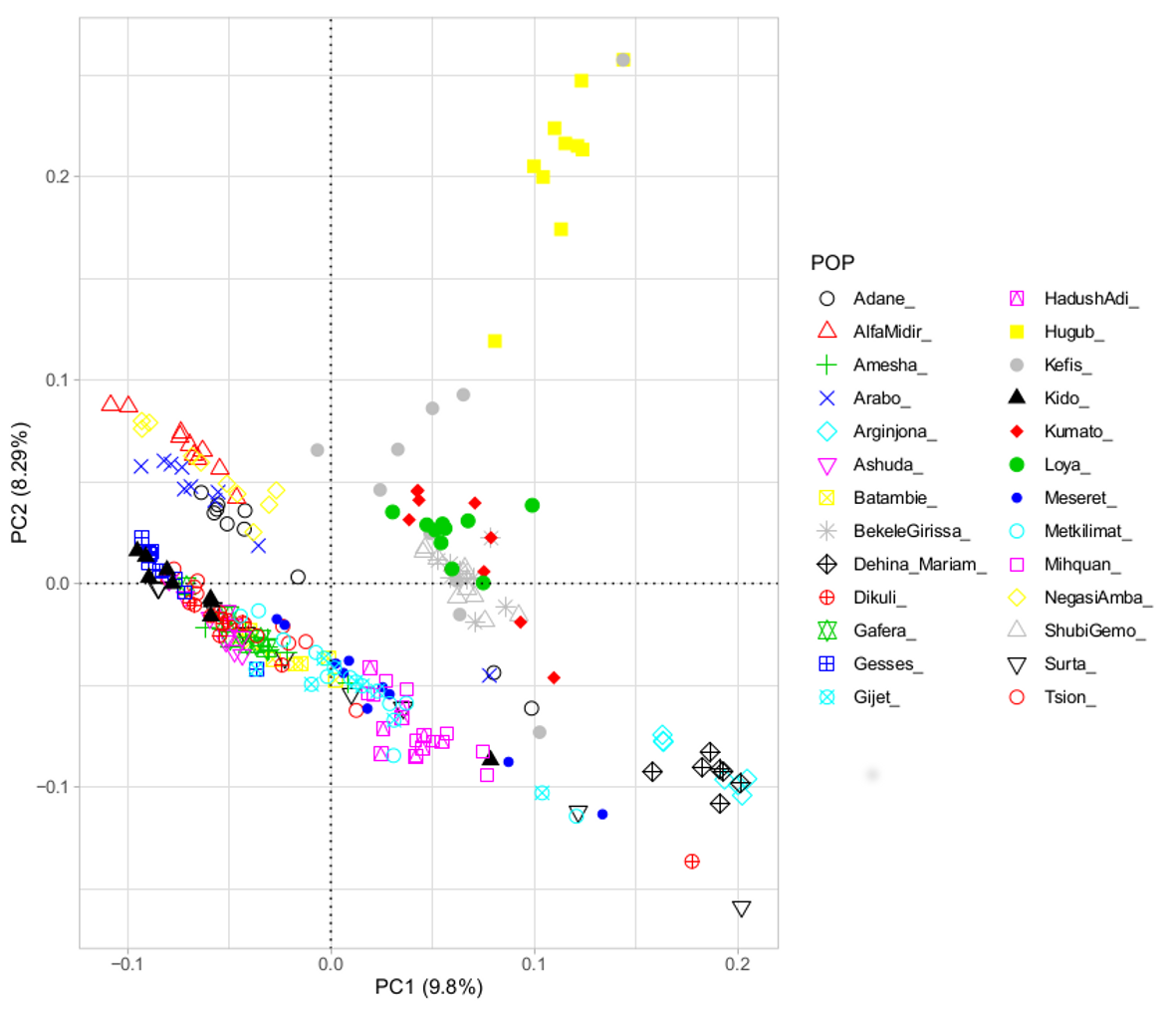

**Principle coordinate analysis showing the clustering of samples by autosomal SNPs. Samples are labelled by the region in which the sample was collected.**

**Figure S8:**

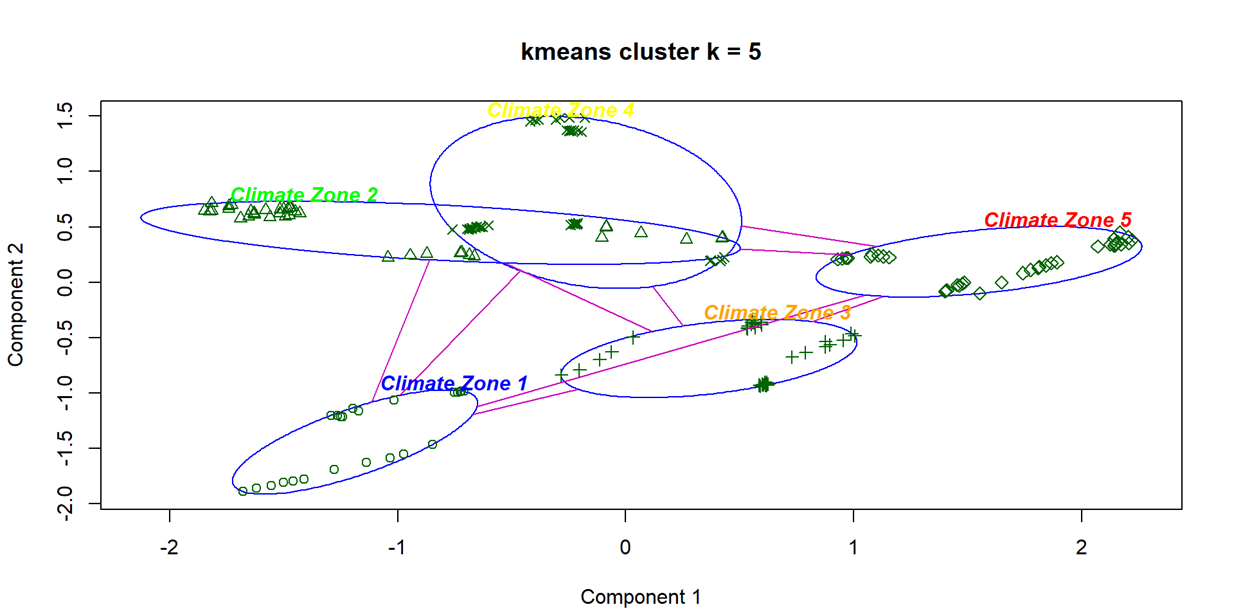

**Five climate zones, clustered using Kmeans according to annual temperature, annual precipitation and precipitation of the driest quarter of the sampling location between 1970 - 2000. Components 1 and 2 explain 85.1% of sampling site variation.**

**Table S1: Diversity of CAZymes in strain level MAGs**

| CAZyme module | Abbreviation | Module function | Families present in strain level MAGs |
| --- | --- | --- | --- |
| Auxiliary activities | AA | Redox enzymes acting in conjunction with CAZymes | 7 |
| Carbohydrate esterases | CE | Hydrolysis of carbohydrate esters | 16 |
| Carbohydrate-binding modules | CBM | Adhesion to carbohydrates | 50 |
| Glycoside hydrolases | GH | Hydrolysis/rearrangement of glycosidic bonds | 126 |
| Glycosyltransferases | GT | Formation of glycosidic bonds | 62 |
| Polysaccharide lyases | PL | Non-hydrolytic cleavage of glycosidic bonds | 24 |
